# Trait selection strategy in multi-trait GWAS: Boosting SNPs discoverability

**DOI:** 10.1101/2023.10.27.564319

**Authors:** Yuka Suzuki, Hervé Ménager, Bryan Brancotte, Raphaël Vernet, Cyril Nerin, Christophe Boetto, Antoine Auvergne, Christophe Linhard, Rachel Torchet, Pierre Lechat, Lucie Troubat, Michael H. Cho, Emmanuelle Bouzigon, Hugues Aschard, Hanna Julienne

## Abstract

Since the first Genome-Wide Association Studies (GWAS), thousands of variant-trait associations have been discovered. However, the sample size required to detect additional variants using standard univariate association screening is increasingly prohibitive. Multi-trait GWAS offers a relevant alternative: it can improve statistical power and lead to new insights about gene function and the joint genetic architecture of human phenotypes. Although many methodological hurdles of multi-trait testing have been discussed, the strategy to select trait, among overwhelming possibilities, has been overlooked. In this study, we conducted extensive multi-trait tests using JASS (Joint Analysis of Summary Statistics) and assessed which genetic features of the analysed sets were associated with an increased detection of variants as compared to univariate screening. Our analyses identified multiple factors associated with the gain in the association detection in multi-trait tests. Together, these factors of the analysed sets are predictive of the gain of the multi-trait test (Pearson’s ρ equal to 0.43 between the observed and predicted gain, *P* < 1.6 × 10^-60^). Applying an alternative multi-trait approach (MTAG, multi-trait analysis of GWAS), we found that in most scenarios but particularly those with larger numbers of traits, JASS outperformed MTAG. Finally, we benchmark several strategies to select set of traits including the prevalent strategy of selecting clinically similar traits, which systematically underperformed selecting clinically heterogenous traits or selecting sets that issued from our data-driven models. This work provides a unique picture of the determinant of multi-trait GWAS statistical power and outline practical strategies for multi-trait testing.

## Introduction

Despite a continuous increase of the sample size of Genome-Wide Association Studies (GWAS), many genetic variants underlying human complex traits and diseases remain undetected. To increase statistical power and detection of associations at low cost, investigators have developed various multi-trait approaches based on GWAS summary statistics^1–6^. Although other factors might also be involved, few studies investigated how the choice of phenotypes impacts the gain in the power of multi-trait approaches. In the standard univariate GWAS, statistical power mostly depends on minor allele frequency, sample size, the size of genetic effect, and polygenicity (the number of causal variants)^7^. In the multi-trait GWAS, it additionally depends on complex characteristics of the set of traits: their shared aetiology, their genetic correlation, and the number of traits in the set. As previously described, the increased power of multi-trait test partly comes from adjusting for the correlation across GWAS studies due to sample overlaps and genetic relationships across phenotypes^4,8,9^. Previous works explored this question using simulated data^8,9^. However, simulations are limited in their scope, and more studies are needed to better characterise scenarios increasing the gain of multi-trait tests.

Here we empirically examined how trait characteristics impact the statistical power of multi-trait GWAS. We performed our analyses on 72 curated GWAS summary statistics and analysed the impact of 11 genetic features describing both individual and collective characteristics of sets of GWAS. Among available multi-trait methods, we used a standard *k*-degree of freedom joint test (omnibus test) of *k* GWAS implemented in JASS^2^ as our primary analysis. The JASS package and its associated tools solve all practical issues commonly encountered in multi-trait GWAS analyses, including missing data and computational efficiency, allowing for a large-scale power analysis on real data. We inspected the association between these genetic features and the gain in the association detection in multi-trait GWAS over univariate GWAS. We then assessed how well these genetic features predict the gain. We further compared JASS and MTAG, an alternative multi-trait approach, in terms of the impact of the trait selection strategy on the gain of multi-trait GWAS.

## Results

### Study Overview

We conducted a series of analyses to identify features that influence the gain in association detection of multi-trait GWAS relative to univariate GWAS. The key principles and main steps of the study are depicted in **Figure 1**. We used 72 curated GWAS summary statistics spanning a range of clinical domains (**Table S1**) and considered 11 features describing the univariate and multivariate genetic architecture of the phenotypes. Five of them characterise single GWAS: mean genetic effect size (MES), polygenicity, effective sample size (*N*_*eff*_), linear additive heritability of common variants (*h*^2^), the proportion of uncaptured linear additive heritability of common variants (%*h*^2^_GWAS_). The remaining six features characterise sets of GWAS. It includes the number of GWAS analysed jointly, and five metrics related to the genetic (Σ_)_, **Table S2** and **Fig. S1**) and residual (Σ_*r*_, **Table S3** and **Fig. S2**) covariance matrices, where the latter represents the covariance between the *Z*-score statistics of two GWAS due to sample overlap and phenotypic correlation. Those five features are the mean of the off-diagonal terms denoted as Σ^’^ and Σ^’^_*r*_; the conditional numbers of Σ_)_ and Σ_*r*_ denoted as Σ_g_ and Σ_*r*_, which we used as a measure of multi-collinearity; and the average difference between the two matrices Δ_+_. Δ_+_ is expected to be a key driver of the power of multi-trait test^8,9^ (**Supplementary note 1, 2, Fig. S3**).

**Figure 1.**
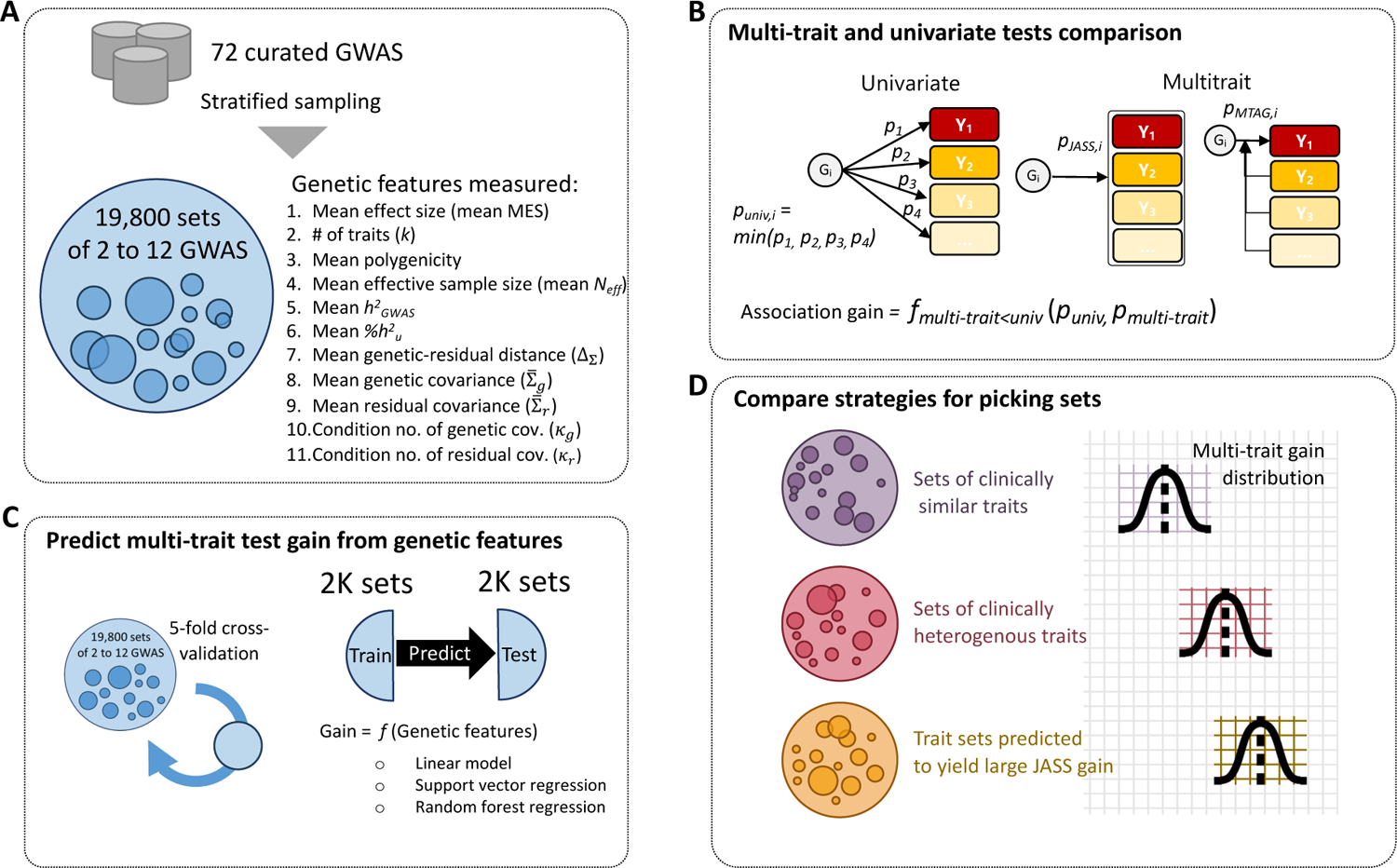
Study overview We conducted a power analysis on real data to understand in which setting a standard multi-trait test—the omnibus test—outperforms univariate GWAS. A) To assemble our real data, we curated 72 GWAS summary statistics and formed about 20k sets of traits by random sampling. Each set of traits was characterised by assessing key genetic features such as polygenicity, mean effect size (MES), and heritability. (B) For each set of traits, we ran omnibus test using JASS and computed the association gain compared to the univariate test. We defined this association gain as the number of LD-independent loci where the omnibus yields a smaller *P*-value than the univariate test. We repeated the analysis using MTAG, a popular multi-trait approach. (C) To investigate which genetic features (highlighted in A) explain the JASS (omnibus) association gain, we applied statistical models to predict the gain as a function of genetic features. Several models were benchmarked to optimise prediction performances. (D) To suggest a practical strategy for selecting traits that yield a large association gain, we compared the performance of JASS on trait sets that are clinically similar, clinically heterogenous, and that were predicted to have a large gain by the predictive model highlighted in (C).

Given the 72 traits, there are over 4.7 × 10^21^ possible sets of two to 72 traits. In this study, we used 19,266 unique random sets of two to 12 GWAS drawn using a stratified sampling conditional on mean effect size, heritability and the number of traits to maximize the range of the genetic features studied (see **Material and methods, Supplementary note 3**). For each set, we derived the average of the single GWAS feature (polygenicity, MES, *N_eff_*, *h^2^_GWAS_*, and *%h^2^_u_*) and the six set features (Σ^’^_)_, Σ^’^_*r*_, Σ_)_, Σ_*r*_, Δ_+_, and the number of GWAS selected). Note that we used the log of polygenicity, MES, Σ^’^_g_, Σ^’^_*r*_, Σ_)_, Σ_*r*_, and Δ_+_ when investigating their association with other variables due to their skewed distributions (**Fig. S4**). Then, we compared multi-trait and univariate GWAS results based on the minimum *P*-value of the multi-trait test across all variants within LD-independent loci (*P*_*JASS*_) and the minimum *P*-value of univariate tests across all variants and all GWAS within the same loci (*P*_*un*i_). We used two metrics: i) the proportion of significant loci found associated at genome-wide level (*P* < 5 × 10^-8^) by JASS and missed by the univariate GWAS, and ii) the fraction of loci where *P*_*JASS*_ was smaller than *P*_*un*i_ ^(*f*^_*JASS*/*un*i*v*_^).^

We first describe the distribution of the genetic features and their correlation. Second, we applied JASS to each set and compared multi-trait and univariate GWAS association results (**Fig. 1B**). Third, we build a predictive model of the association gain using a five-fold cross-validation approach (**Fig. 1C**). For each cross-validation, we pulled a training set of 1,980 out of the 19,266 unique random sets and conducted a regression analysis to estimate the contribution of a subset of genetic features of trait sets using a joint model of the features. We measured the correlation between observed and predicted gain in an independent validation dataset. Fourth, we compared the association gain from JASS against MTAG^4^, a popular multi-trait approach that leverages genetic correlation across traits to boost statistical power. Finally, we compare common trait selection strategies—e.g. choosing clinically homogenous or heterogeneous traits—in their impacts on the association gain to evaluate our model prediction and provide a practical strategy in multi-trait test (**Fig. 1D**).

### Distribution of the genetic features across GWAS trait sets

The individual genetic features of the 72 studied traits were distributed as follows. The sample sizes ranged from 5,318 to 697,828 with a median of 85,559. Heritability (*h*^2^_GWAS_) ranged from 1% to 48% with a median at 10% (**Fig. 2A**) and was consistent across the software used for its estimation (**Fig. S5**). Polygenicity and MES were highly variable: Polygenicity (i.e. the estimated number of causal variants) ranged from 6.9 (Estimated Glomerular Filtration Rate from Cystatin C) to 570,102 (variability sleep duration) with a median of 1,110, and MES ranged from 4.48 × 10^-8^ (variability sleep duration) to 3.3 × 10^-3^ (Fasting proinsulin) with a median of 1.0 × 10^-4^. The distribution of the average these parameters across the 19,266 random sets is presented in **Figure 2B** along with the GWAS set features. The latter metrics ranged (in 25-75 percentiles) as −2.04 to −1.79 for log_12_ Σ^’^_)_, −1.96 to −1.44 for log_12_ Σ^’^_*r*_, 0.36 to 0.78 for log_12_ Σ_)_, 0.05 to 0.32 for log_12_ Σ_*r*_, and −1.11 to −0.89 for log_12_ Δ_+_. In particular, the variability in log_12_ Δ_+_ was limited (given the theoretical upper bound of log_12_ Δ_+_ is log_12_ 2 ≅ 0.3).

**Figure 2.**
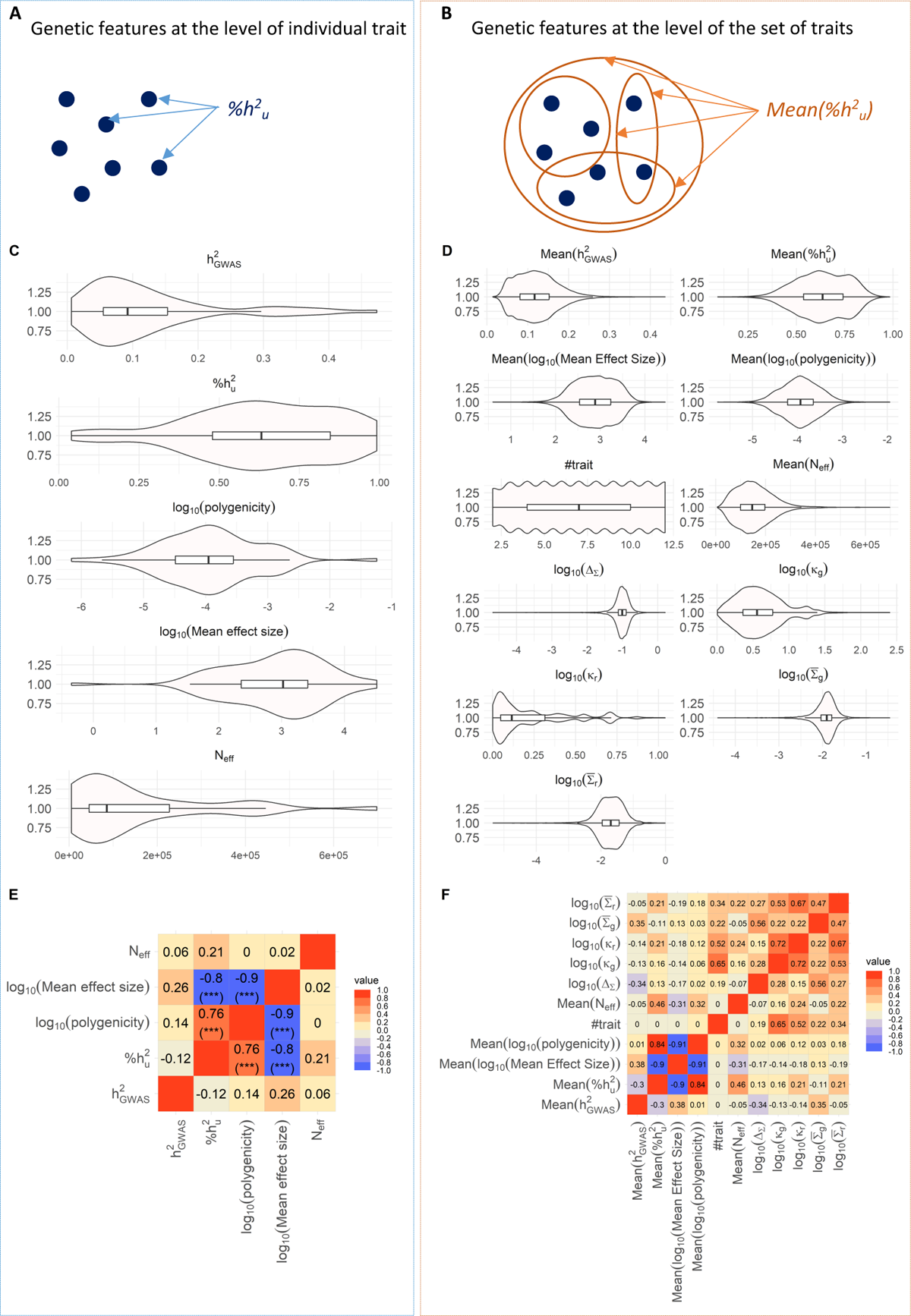
Genetic features characteristics derived from 72 traits and 19,266 random trait sets. Visualization of the investigated features and their relation at the level of individual trait (panels A, C, and E) and at the level of set of traits (panels B, D, and F). A) Schematic of a genetic feature derived at the level of an individual trait. B) Schematic of a genetic feature derived at the level of a set of traits. C) Violin plots representing the distribution across the 72 summary statistics of heritability, polygenicity, MES and sample size of the study. D) Violin plots representing the distribution across the 19,266 sets of traits of the 11 genetic features derived for each trait set. E) Pearson correlation among polygenicity, MES, *h^2^_GWAS_*, *%h^2^_u_*, and sample size across 72 traits. F) Pearson correlation among the 11 features across 19,266 trait sets. (q-value annotation: *** < 10^-3^, ** <10^-2^,*< 5×10^-2^)

We measured the correlation across features at the trait and set levels (**Fig. 2C, 2D**). The proportion of uncaptured linear additive heritability of common variants (%*h*^2^_GWAS_) was strongly associated with log_12_ polygenicity (ρ = 0.76, *P* = 8.8 × 10^-15^) and with log_12_ MES (ρ = −0.8, *P* = 3.5 × 10^-17^) (**Fig. 2C, S6**), in agreement with previous reports showing that univariate GWAS performs better for traits with a larger MES and a smaller polygenicity^7^. As expected, the means of individual trait feature were correlated in the same way across sets as across traits (e.g. mean %*h*^2^_GWAS_and mean polygenicity are highly correlated as are %*h*^2^and polygenicity, **Fig. 2D**). Metrics of genetic and covariance matrices were positively correlated to each other, i.e. log_12_ Σ^’^ with log_12_ Σ^’^_*r*_and log_12_ Δ_+_, log_12_ Σ_)_ with log_12_ Σ_*r*_, and log_12_ Σ^’^_*r*_with log_12_ Σ_*r*_ and log_12_ Σ_)_, which further correlated with the number of traits.

### Multi-trait versus univariate GWAS across 19,266 random sets

We applied JASS to all 19,266 sets and quantified the gain in association detection of the multi-trait against univariate test. On average, in a set, JASS detected 26 new loci while 285 were previously associated with the univariate tests, i.e. a 1.1-fold increase in the total number of loci detected (**Fig. S7A**). JASS gain was maximal for sets with a relatively low number (< 300 loci) of previously detected loci (**Fig. S7B**). JASS detected at least one new association in 98% of the trait sets (18,787 sets, **Fig. S7C**), and 508,829 new associations in total (note that these can be overlapping loci). These numbers are obtained when applying JASS on variants with beta coefficients available for all traits in the set. When analysing all variants including those with missing values as allowed by JASS (see **Material and methods**), about 1.4 times more new associations were detected (693,382 new associations in total by JASS including variants with missing values, **Fig. S7D**).

We assessed the marginal relationship between genetic features and the number of loci detected by the univariate and multi-trait GWAS tests, and the association gain of multi-trait test. **Figure 3** presents the correlation between each feature and the number of univariate and multi-trait GWAS associated loci. As expected, the number of univariate associated loci was positively correlated with the number of traits, mean *N_eff_*, the mean *h^2^_GWAS_* and mean MES, and negatively correlated with mean uncaptured linear additive heritability (mean %*h*^2^). Multi-trait GWAS gain over univariate GWAS was positively associated with mean polygenicity and mean %*h*^2^_GWAS_while negatively associated with mean MES. Mean *h²_GWAS_* show opposite effect, being slightly positively associated with gain, but negatively associated with gain. Overall, this suggests that the multi-trait test can be highly complementary to the univariate test, performing better in situations where the univariate tests display low power. We noted in a recent study^10^ that a high multicollinearity of the matrix underlying the null hypothesis (Σ_*r*_) can lead to a lack of robustness of the omnibus test^10^. We checked how the condition number Σ_*r*_was related to JASS gain (**Fig. 3B**, **Fig. S8**). The condition number stayed in a reasonable range for 99% of sets (min=1, max=11). The rest (1%) of sets were flagged for having singular residual matrices. Furthermore, note that the trait sets contained overlapping traits and are therefore not fully independent from each other. To robustly assess the impacts of genetic features on the multi-trait gain by addressing this issue, below we conducted regression analyses with a cross validation scheme establishing independence between training and validation data (**Fig. S9**).

**Figure 3.**
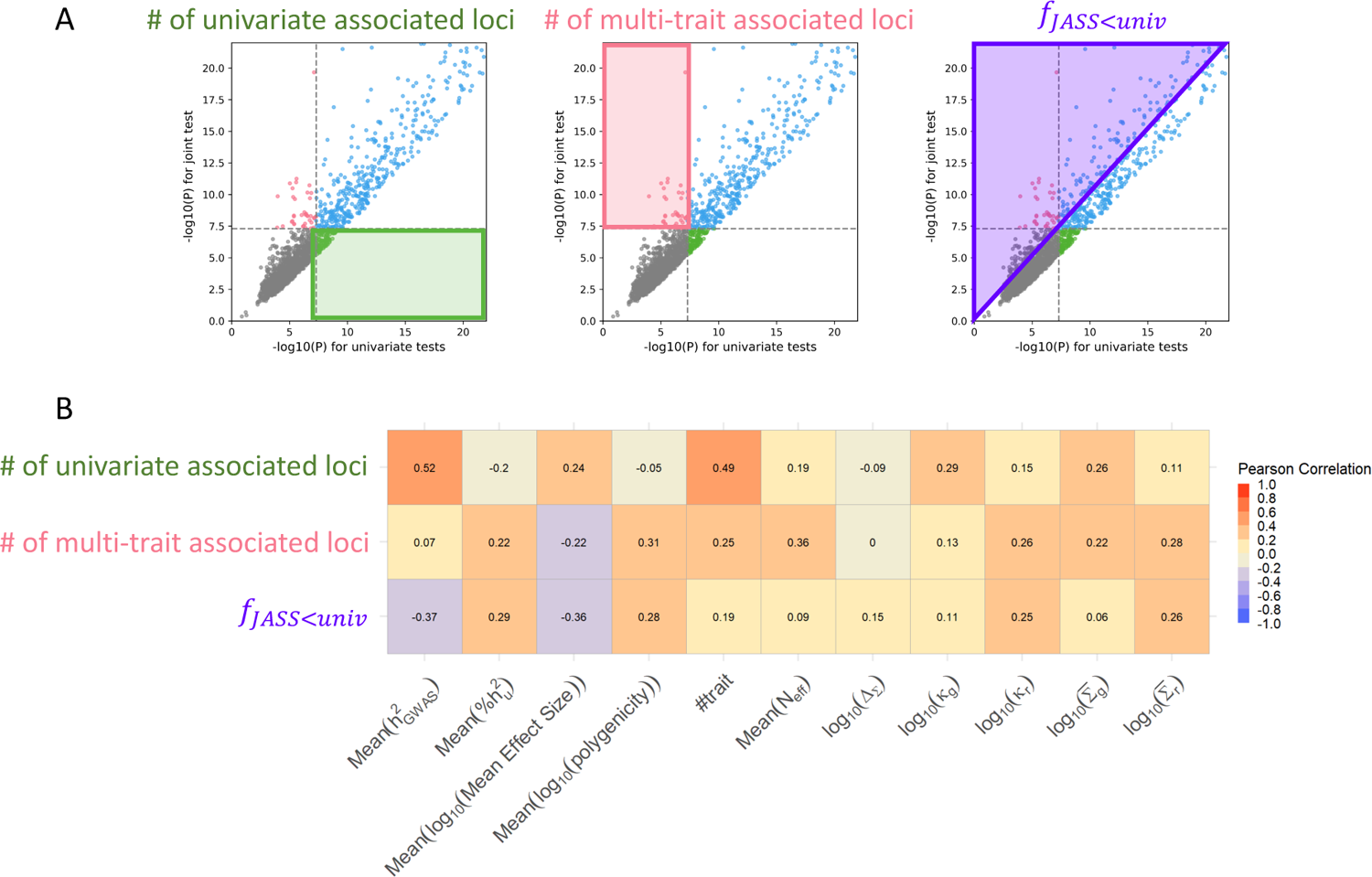
Determinant of JASS gain across trait sets. A) Illustration of the three metrics to assess univariate and multi-trait GWAS outcomes. On a quadrant plot representing the *P*-value of the multi-trait test with respect to the *P*-value of the univariate test, the following areas represent regions where: (green) only the univariate test is significant, (pink) only the multi-trait test is significant, (purple) the multi-trait test is more significant than the univariate test. B) Heatmap of the Pearson correlation between the number of univariate association loci, the number of new association loci detected by JASS, the association gain of JASS (*f_JASS<univ_*) and the 11 genetic features across 19,266 trait sets.

### Predicting multi-trait test gain from genetic features

We investigated the predictive power of the association gain (*f*_*JASS*/*un*i*v*_) from a joint modelling of the genetic features (**Fig. 4**). Note that some features are almost linear combinations of other ones (e.g. Δ_+_ is proportional to the difference between Σ^’^ and Σ^’^_*r*_). To avoid extreme collinearity and ensure parsimony, we selected six moderately correlated features: the number of traits, log_12_ Δ_+_, log_12_Σ^’^_)_, mean *N_eff_*, mean *h²_GWAS_*, and mean %*h*^2^(see **Material and methods**). We favoured mean%*h*^2^_GWAS_over mean MES and mean polygenicity since mean %*h*^2^_GWAS_captures both log_10_(MES) andlog_10_(polygenicity) (**Fig. 2**). We assessed performances using a five-fold cross validation (CV). For each CV the model parameters were derived on a training data, and the prediction accuracy was derived in an independent data (**Material and methods, Fig. S9**). All six features were highly associated to the multi-trait gain (**Table 1**). Mean %*h*^2^_GWAS_was overall positively associated to multi-trait gain, whereas mean *h²_GWAS_* was negatively associated. Overall, the association gain of multi-trait test, more specifically the omnibus test, seems driven by genetic correlation, polygenicity, and the number of traits. The predicted gain was significantly correlated with the observed gain on validation data (median Pearson ρ=0.43, *P* < 1.6 × 10^-60^, **Fig. S10**).

**Figure 4.**
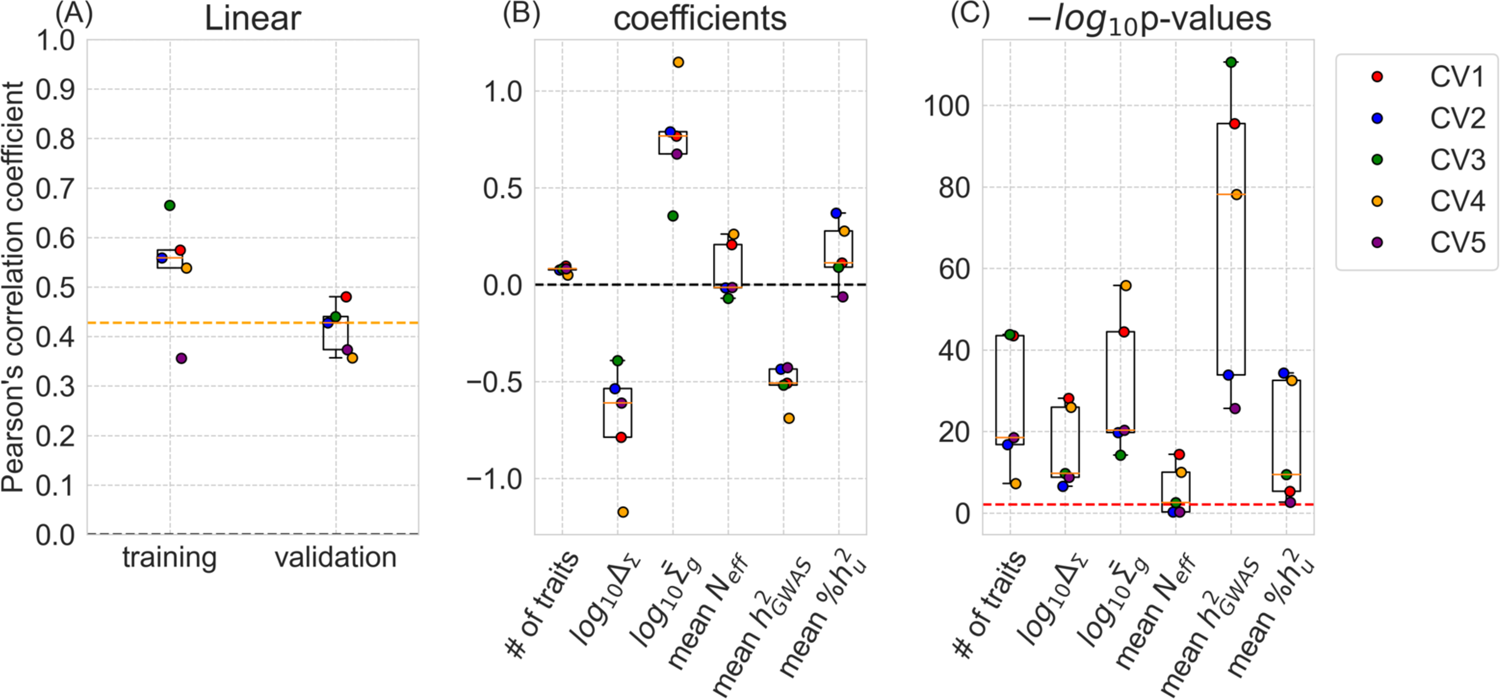
Model prediction power and feature contributions. (A) Boxplots of the prediction power across the five-fold cross validations (CV) of the multivariate linear regression model measured as the Pearson’s correlation coefficient between the predicted and observed gain. The performance of each CV is represented as a coloured dot. Orange dashed line: median correlation coefficient between the predicted and observed gain in the validation data. (B, C) The boxplots show the coefficients and -log_10_(*P*-values) of the six features in the regression model across five-fold cross validations using each corresponding training data. Red dashed line: Bonferroni corrected nominal significance threshold.

**Table 1.** Coefficients of the multivariate linear regression models from five-fold cross validations. Mean genetic residual distance and mean genetic covariance were log_10_ transformed. All the features were scaled using *MinMaxScaler* in *scikit-learn*^37^.

|  | CV1 | CV2 | CV3 | CV4 | CV5 | Mean |
| --- | --- | --- | --- | --- | --- | --- |
| # of traits | 0.096<br>(p= 3.447x10<sup>-44</sup>) | 0.077<br>(p= 1.592x10<sup>-17</sup>) | 0.082<br>(p= 1.638x10<sup>-44</sup>) | 0.049<br>(p= 6.148x10<sup>-8</sup>) | 0.083<br>(p= 3.071x10<sup>-19</sup>) | 0.077<br>(std=0.017) |
| Mean $\log_{10} \Delta_{\Sigma}$ | -0.787<br>(p= 8.338x10<sup>-29</sup>) | -0.536<br>(p= 8.338x10<sup>-29</sup>) | -0.393<br>(p= 2.242x10<sup>-10</sup>) | -1.174<br>(p= 1.086x10<sup>-26</sup>) | -0.610<br>(p= 1.947x10<sup>-09</sup>) | -0.700<br>(std=0.301) |
| Mean $\log_{10} \bar{\Sigma}_g$ | 0.767<br>(p= 3.813x10<sup>-45</sup>) | 0.789<br>(p= 2.055x10<sup>-20</sup>) | 0.355<br>(p= 5.990x10<sup>-15</sup>) | 1.149<br>(p= 1.604x10<sup>-56</sup>) | 0.675<br>(p= 5.636x10<sup>-21</sup>) | 0.747<br>(std=0.284) |
| Mean $N_{eff}$ | 0.206<br>(p= 4.342x10<sup>-15</sup>) | -0.016<br>(p= 0.554) | -0.071<br>(p= 0.003) | 0.260<br>(p= 1.099x10<sup>-10</sup>) | -0.015<br>(p= 0.598) | 0.073<br>(std=0.149) |
| Mean $h^2_{GWAS}$ | -0.509<br>(p= 2.937x10<sup>-96</sup>) | -0.437<br>(p= 1.455x10<sup>-34</sup>) | -0.518<br>(p= 2.216x10<sup>-111</sup>) | -0.690<br>(p= 6.670x10<sup>-79</sup>) | -0.429<br>(p= 2.225x10<sup>-26</sup>) | -0.516<br>(std=0.105) |
| Mean $\%h^2_u$ | 0.112<br>(p= 5.158x10<sup>-6</sup>) | 0.370<br>(p= 4.353x10<sup>-35</sup>) | 0.091<br>(p= 3.877x10<sup>-10</sup>) | 0.277<br>(p= 3.022x10<sup>-33</sup>) | -0.064<br>(p= 0.002) | 0.157<br>(std=0.169) |

We conducted a series of sensitivity analyses to explore further performances. First, to interpret this model behaviour from the perspective of polygenicity and MES, we fitted two models replacing mean %*h*^2^_GWAS_with mean log_10_(polygenicity) and mean log_10_(MES) in turn (**Fig. S11**). In these models, mean log_10_(polygenicity) had a positive contribution to the gain, whereas mean log_10_(MES) had a negative contribution. In other words, multi-trait tests likely detect new associations in settings where univariate tests perform poorly, confirming the correlation analysis (**Fig. 3**). Additionally, the model suggested the number of traits and log_12_ Σ^’^ further enhance multi-trait test gain. Contrary to previous observation on simulation studies, log_12_ Δ_+_ likely diminishes the multi-trait test gain (see **Supplementary note 4** for a hypothetical explanation). Second, we also considered two nonlinear models for comparison purposes, support vector regression and random forest regression. As showed in **Figure S12**, these two models appear to outperform the multivariate linear model on the training datasets. However, they performed similarly or worse on the test dataset, suggesting a strong overfitting in the training data. Overall, we did not find any benefit in using these more complex models.

### Comparison of JASS versus MTAG

We repeated the association screening and the prediction analysis using MTAG (Multi-Trait Analysis of GWAS), a popular multi-trait approach leveraging genetic correlation among closely related traits to inform GWAS screening. Regarding the association screening, MTAG detected 153,061 new association regions in total across the sets (30% of the number of new associations detected by JASS). On average, MTAG detected eight new association loci per set and at least one new association locus on 63.3% of sets (12,195 out of 19,266) compared to 98% of sets for JASS. In 93% of all the trait sets, MTAG detected fewer associations than JASS did (**Fig. 5A**). The performance difference further increased when applying JASS also on variants with missing values (in 96% of trait sets MTAG detected fewer associations). Despite these discrepancies in the number of association loci, the number of new association loci in MTAG and JASS were strongly correlated (Pearson ρ = 0.72, *P* < 2.2 × 10^-308^), and as well as the *P*-values of MTAG and JASS for the same loci (Pearson ρ =0.75, *P* < 2.2 × 10^-308^). This concordance suggests common determinants for statistical power between the two methods. We next fitted a multivariate linear model to predict MTAG gain from the same six genetic features and training data used for JASS (**Fig. S13**). The most notable difference from JASS was that the number of traits in the set was, consistently across cross-validation folds, negatively associated with the gain of MTAG. Indeed, JASS particularly outperformed MTAG on larger set of traits (**Fig. 5B**).

**Figure 5.**
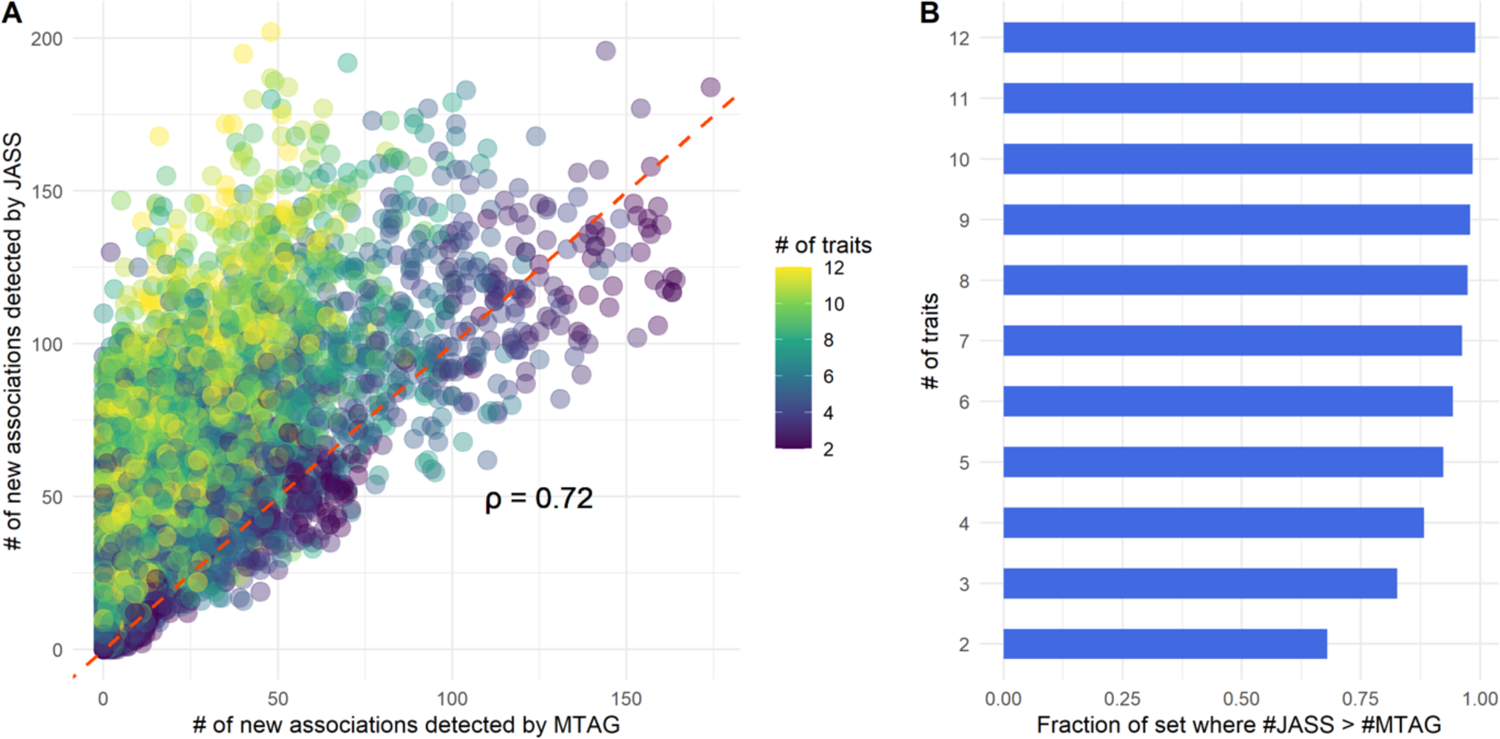
Comparison of MTAG with JASS. A) The number of new association loci found by JASS with respect to the number of new association loci found MTAG across all the trait sets. Each dot represents a set of traits. Dot colours represent the number of traits in the set. B) Fraction of sets where the number of new association loci detected by JASS was superior to the number of associations detected by MTAG stratified by the number of traits in the set.

To test whether the pronounced gain of JASS compared with MTAG on large sets was mostly due to multiple testing correction—as MTAG performs one test by trait present in the set of traits, multiple testing correction is applied to MTAG *P*-values (**Material and methods**)—we compared the number of associations detected by JASS and by MTAG without correction (**Fig. S14**). In the absence of multiple testing correction, MTAG type 1 error expectedly increased with the number of traits in the set (e.g. genomic inflation factor λ was 1.3, 1.66, 1.72, and 1.77 for four cases with 2, 5, 9, and 12 traits). Despite the increased *P*-value inflation for large sets, MTAG detected fewer associations overall (**Fig. S14A**) than JASS on most of the sets (in 71% and 83% of sets when excluding and including variants with partly unavailable summary statistics, respectively). The performance of JASS and MTAG were most similar in small trait sets (**Fig. S14B**), while JASS was particularly advantageous when analysing large trait set especially when allowing for variants with summary statistics partly unavailable.

### An informed strategy for trait set selection in multi-trait GWAS

A common and seemingly sound strategy when conducting multi-trait analyse is to use closely related traits. This choice is partly driven by investigators’ interest in delineating the shared genetic aetiology between a disease and closely related phenotypes. This might also arise from the intuitive idea that closely related phenotypes share a fair amount of genetic aetiology that the multi-test could leverage. However, its impact on statistical power has not been evaluated. To advise investigators on the best strategy to compose sets, we compared multi-trait gain and the number of new association loci obtained using this clinically-driven strategy (referred as “*homogenous*”, see **Material and methods**) to three alternatives: i) including GWAS from two to four clinical groups (noted “*low heterogeneity*”), ii) including GWAS from five clinical groups or more (noted “*high heterogeneity*”), and iii) a data-driven approach based on the linear regression predicted gain (**Material and methods**). The data-driven strategy had a higher gain and larger number of new association loci by JASS compared to the other strategies (**Fig. 6A and 6B**). The gain and the number of new association loci increased systematically with the clinical heterogeneity of the traits, and the increases were statistically significant for most pairs of trait selection strategies compared, especially when comparing the data-driven approach with the other approaches (*P* < 4.4 × 10^-8^ for gain and *P* < 6.7 × 10^-7^ for the number of new association loci). The average number of new association loci detected in validation data equalled 14, 20, 32, and 61 for homogeneous, low heterogeneity, high heterogeneity, and data-driven sets, respectively.

**Figure 6.**
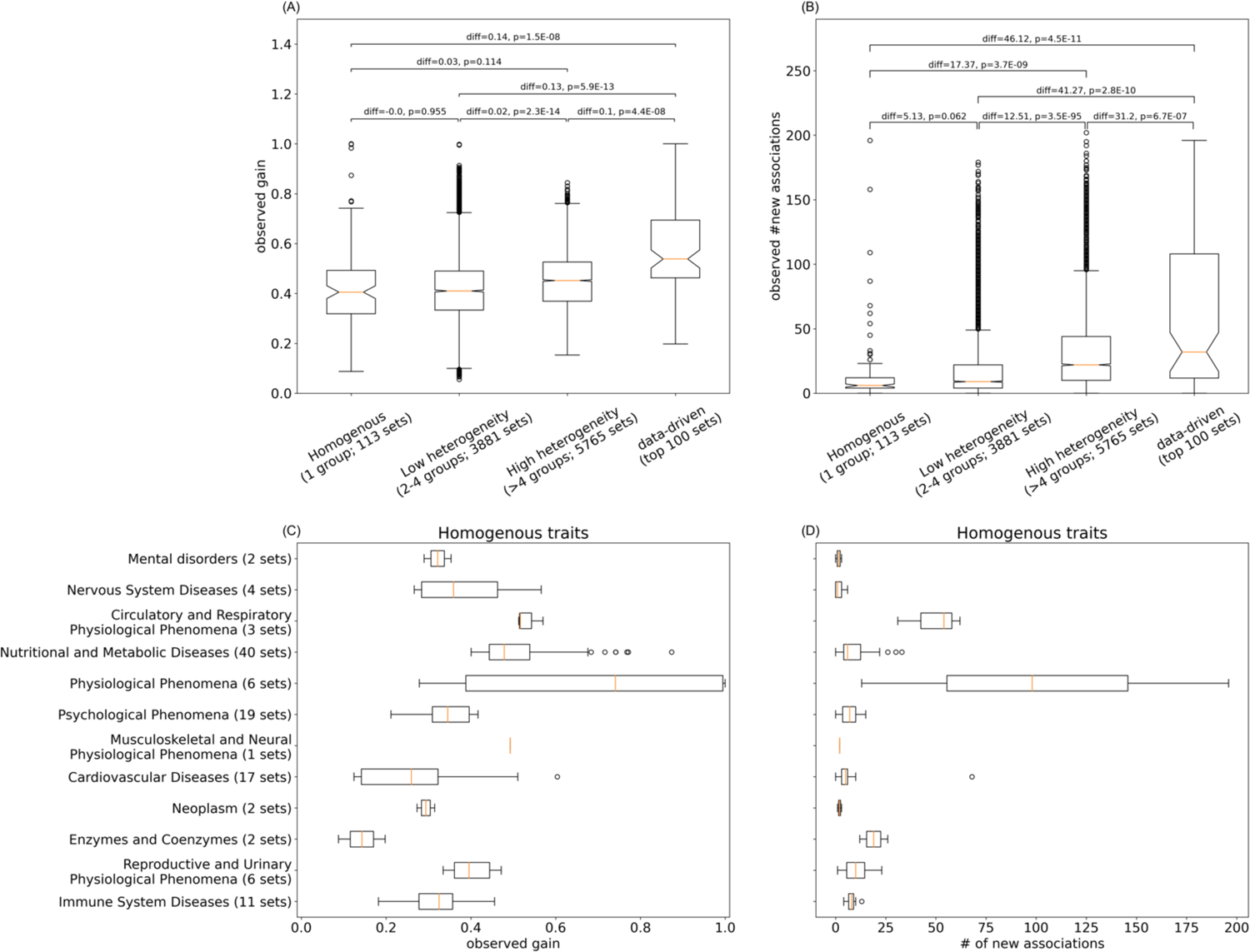
Comparison between clinical and data-driven trait sampling methods. (A, B) Distribution of the gain and the number of new association loci for trait sets selected by four trait selection strategies from the validation data. *P*-values are from the two-sided Welch’s *t*-test. Differences in mean values in each pair compared in the test (right – left categories in the order shown on the x-axis) are also shown. Note we used a pair of a training data and a validation data to ensure independence for the test. The numbers under the labels on the x-axis indicate the number of trait sets from each strategy. The observed JASS gain and the number of new association loci are shown on the y-axis. (C,D) The observed gain and number of new association loci detected for trait sets of the ‘homogenous’ category, visualised per clinical grouping.

We further decomposed the gain of the homogenous sets by clinical groups (**Fig. 6C, 6D**). Two groups consistently yielded more new association loci than others, namely: ‘Circulatory and Respiratory Physiological Phenomena’ and ‘Physiological Phenomena.’ The first group was composed of spirometry traits and asthma, characterised by a large sample size, substantial genetic correlation, and moderate heritability: a favourable setting for JASS. The second group contained anthropometric traits (Height, BMI, Hip circumference, waist circumference, and Waist hip ratio). We next investigated whether multi-trait gain was associated with specific traits and derived the fold enrichment for traits between the top 10% yielding sets (corresponding to >169 new associated loci by JASS) and the least 10% yielding sets. BMI and hip circumference were amongst the top traits (**Fig. S15**). Notably, we found that high yielding trait sets (i.e., those that yielded 169 or more new association regions by JASS) all contained BMI. In contrast, out of the remaining trait sets, only 9% of the trait sets contained BMI. Other traits enriched in top sets included spirometry and to a lesser extend mental disorders, arterial pressure, and sleep patterns. BMI genetics is increasingly recognized to be a complex entanglement of metabolic and behavioural factors^11^, and suggests that complex traits that reflect multiple biologic processes may benefit more from multi-trait analysis.

We additionally compared the multi-trait gain and the number of new association loci by MTAG across the four groups of trait sets. The results were rather opposite to what we observed above with JASS: MTAG gain increased as the trait sets became more homogenous (**Fig. S17A**, *P* < 1.4 × 10^-7^), and the number of new association loci was greater for trait sets of low heterogeneity than high heterogeneity (**Fig. S17B**, *P* = 2.7 × 10^-6^). We then tested whether MTAG outperforms JASS on homogenous trait sets (**Fig. S17C,D**) and found that there was little difference between the two (*P*=0.084 for gain, *P*=0.063 for the number of new association loci). We further tested whether MTAG outperforms JASS on homogenous trait sets of certain clinical groups (**Fig. S17E,F**). JASS significantly outperformed on ‘Immune System Diseases’ and ‘Psychological Phenomena’ (‘Immune System Diseases’: *P* = 8.2 × 10^-4^ for gain and *P* = 6.7 × 10^-7^ for the number of new association loci, ‘Psychological Phenomena’: *P* = 2.5 × 10^-5^ for the number of new association loci). In contrast, MTAG outperformed on ‘Musculoskeletal and Neural Physiological Phenomena’ (which contains only one trait set).

While JASS type 1 error in JASS is properly calibrated, this large number of new associations could be spurious, and irrelevant to biology. To ensure that new associations detected by JASS are relevant, we focussed on BMI and tested if we could predict novel associations observed in a larger study (sample size= 683,365 ^12^) from multi-trait GWAS applied on the BMI study in the present analysis (sample size= 339,224 ^13^, **Table S1**). Across the 1,776 sets containing BMI, JASS detected 1,167 new associations (after Bonferroni correction, **Material and methods**) of which 537 corresponded to a new association in the larger BMI GWAS. 86 associations of the larger GWAS were missed by JASS. Hence, JASS was able to flag loci with a high recall (0.86, probability of new association detected in the larger GWAS to be detected by JASS) but a moderate specificity (0.46, probability of a detected loci to be associated in the larger GWAS). This can be explained by the generality of the null hypothesis used in JASS which requires only one trait in the set to be significant (not necessarily BMI). To improve specificity, we fitted a logistic regression predicting if a locus would be associated with BMI in the larger GWAS by combining the number of sets where the locus was associated by JASS and the minimum *P*-value across sets (**Fig. S16**). This model reached an AUC of 0.75, an accuracy 0.74, a recall of 46% and a specificity of 75% when applying a standard probability threshold of 0.5. Based on JASS results, we were able to infer a substantial number of loci associated in a GWAS with twice as many samples as the one used in the present study.

## Discussion

This study investigated the genetic features associated with the statistical power of multi-trait GWAS. On average, the power increase relative to the univariate GWAS was substantial: JASS detected new association loci in 98% of 19,266 sets, with an average of 26 new association loci. This power increase appears to be highly associated with the genetic features of trait sets. More specifically, multi-trait gain tends to be higher for sets with 1) a moderate mean heritability (mean *h*^2^_GWAS_), 2) a smaller mean MES, 3) a larger mean polygenicity, 4) a larger genetic covariance across traits, and 5) a smaller distance between the residual and genetic covariance. Although some features, such as increased distance between genetic and residual correlation, have been previously found associated with the gain in association detection, this analysis suggests a negative or negligible gain in real data.

This might be explained by the complex correlation across features, highlighting the importance of considering multiple features jointly or by variations of genetic architecture being narrower in real data than assumed by previous simulation studies. We also found that selecting specific traits such as BMI, and more clinically heterogenous sets, specifically for the omnibus test, can strongly outperform approaches that select clinically homogenous sets. Finally, we investigated the predictive power of a multivariate linear model that could predict trait sets that most likely benefit from the multi-trait test (median Pearson’s ρ=0.43, *P* < 1.6 × 10^-60^), which can be used to astutely select traits to be tested jointly. Our findings provide an approach that can increase the identification of genetic associations using existing GWAS data, with relevance to traits in which genetic signal is scarce. We summarize in **Figure S18** a guideline for selecting the best suited methods according to the feature of their data.

Selecting clinically homogenous traits is the strategy most commonly used^4,8,10,14–19^. In our previous large-scale analysis^8^, an heterogeneous set yielded the largest number of new association as compared to clinically homogenous sets. In this work we further showed that a data-driven strategy is expected to outperform other strategies based on clinical insights. Such sets might capture highly pleiotropic signals hard to detect using univariate GWAS and recommend that investigators compose a heterogeneous set or use our predictive model to build a set of traits.

We compared the results from the standard omnibus test (implemented in JASS) against MTAG, a popular multi-trait GWAS method^20–23^. For the data we used, the omnibus almost systematically outperformed MTAG, with an overall three-fold increase in the number of loci detected. The difference was particularly striking on larger trait sets. MTAG is built on the hypothesis of homogeneous genetic correlation across genetic variants, and therefore is expected to have maximum power when this assumption is valid. Indeed, we observed that more homogenous trait sets yielded larger MTAG gain than heterogenous trait sets (**Fig. S17A,B**). Yet, the omnibus generally performed equally well or better than MTAG even on the homogenous trait sets (**Fig. S17C-F**). In contrast, by construction, the omnibus test allows for substantial heterogeneity, although at the cost of an increase in the degree of freedom. In line with previous work suggesting that genetic correlation might be fairly heterogenous across the genome^24^, this cost appears to be outweighed by the additional flexibility in capturing heterogeneous multi-trait genetic patterns^8^ (**Supplementary notes 1, 2**). Thus, the omnibus test is recommended for general identification of variants that impact phenotypes, while MTAG is suitable for identifying variants associated with specific traits (**Fig. S18**).

Our study has some limitations. First, we focused on commonly measured genetic features, but based on these results, a number of other refined metrics could be used. These include effect size distribution as measured by the alpha parameters^25^. Second, we considered GWAS derived from common diseases and anthropometric traits. Future studies might explore performances using a wider variety of molecular traits, for which GWAS summary statistics are becoming increasingly available. Third, the estimation of the features might be also refined. Here, we used MiXeR^26,27^ to estimate most features. However, we observed a dependency of MES and polygenicity on *N_eff_*. Improving these metrics could improve the overall analysis and interpretations of the results. Fourth, we focused on European ancestry summary statistics. This decision was motivated by the availability of large GWAS and using one ancestry for linkage disequilibrium; however, by doing so, we disposed of many traits with which we could have had a wider variety of genetic features, which might have improved the performance of the predictive model. This focus should not lead the reader to think that multi-trait GWAS is useful only on large sample studies of European ancestry. We actually recently updated the JASS pipeline to run a Multi-ancestry Multi-trait GWAS, which was able to detect 367 new association loci, despite the modest sample size of the non-European cohorts used^10^. Future work might leverage non-European existing^28^ and upcoming biobanks^29^ to investigate the validity of our results for non-European ancestries.

In conclusion, this study provides a first overview of what to expect when applying multi-trait tests to a variety of data and how to maximise new discoveries. These insights can be leveraged to discover genetic variants associated with human complex traits and diseases missed by univariate analysis at no cost. Beyond mapping, JASS used on clinically heterogeneous trait sets might offer a way to understand a shared genetic aetiology among unexpected traits^30^ and contribute to deeper understanding of pleiotropy.

## Acknowledgments

This research was supported by the Agence Nationale pour la Recherche (GenCAST, ANR-20-CE36-0009). This work has been conducted as part of the INCEPTION program (Investissement d’Avenir grant ANR-16-CONV-0005). MHC was supported by R01HL162813, R01HL153248, R01HL149861, and R01HL147148.

## Author contributions

Conceptualization: H.J., H.A.

Methodology: Y.S., H.J., H.A., C.B., A.A., M.H.C.

Software: Y.S., H.J., H.M., B.B., P.L., R.T., C.N., L.T. Validation: Y.S., H.J., H.A., M.H.C., E.B.

Formal analysis: Y.S., H.J. Investigation: Y.S., H.J. Resources: H.A.

Data Curation: H.J., C.L., L.T.

Writing (original draft): Y.S., H.J., H.A.

Writing (review & editing): Y.S., R.V., C.B., A.A., M.H.C., E.B., H.A., H.J. Visualization: Y.S., H.J., H.A.

Supervision: H.J., H.A.

Project administration: H.J., H.A., E.B. Funding acquisition: H.J., H.A., E.B., M.H.C.

## Web resources

JASS https://jass.pasteur.fr/

MeSH Browser https://meshb-prev.nlm.nih.gov/search

## Data and Code availability

https://gitlab.pasteur.fr/statistical-genetics/jass_suite_pipeline

https://gitlab.pasteur.fr/statistical-genetics/jass

https://gitlab.pasteur.fr/statistical-genetics/multitrait_power_traitselection

## Material and Methods

### Database of curated summary statistics

We assembled a database of 72 genome-wide GWAS summary statistics of quantitative traits and diseases conducted in European ancestry population pulled from the GWAS catalogue^31^ and a variety of publicly available meta-analyses. We cleaned, harmonised and imputed each study using our previously developed pipeline^2^. In brief, the process includes the following steps: 1) alignment of each GWAS to the 1000G GRCh37 reference panel^32^, 2) imputation of missing summary statistics using RAISS ^33^, 3) computation of the heritability, genetic and residual covariance matrices, referred further as *h*^2^_GWAS_, Σ_g_ and Σ_r_, using LD-score regression^34^, 4) aggregation of curated GWAS in a unique entry file used as input for JASS. We filtered all GWAS with negative heritability, resulting in a total of 72 traits (**Table S1**). Curated GWAS summary statistics used in the analysis are available on the JASS webserver https://jass.pasteur.fr/.

### Joint test and association gain

Multi-trait analyses were conducted using the omnibus test implemented in the JASS package^2,8^. For a set of *k* GWAS, the omnibus statistics is defined as *T*_omni_ = Z^t^Σ_r_ Z where Z is the vector of *Z*-scores across traits Z = (Z_1_ … Z_*k*_) and Σ_r_ is the residual *Z*-score covariance derived using the LD-score regression. Under the null hypothesis of no association with any of the *k* phenotypes, *T*_omni_ follows a X^2^distribution with *k* degree of freedom. To maximize data usage, the default setting of JASS uses all variants even those with missing association statistics. In this case, JASS returns association *P*-value based on the subset of *Z*-scores available. In contrast, the *–remove-nans* option removes variants with incomplete data. Here, we used *–remove-nans* option as the primary analysis for a better characterisation of trait sets and for a fair comparison with the default setting of MTAG^4^ that do not allow for missing statistics, whereas we also provide some results from the default setting of JASS as an additional information.

All power comparisons were conducted at a locus level. The entire genome was split into a total of 1,703 quasi-independent loci defined based on linkage disequilibrium (LD)-independent blocks, as proposed by Berisa and Pickrell^35^. For both multi-trait and univariate analyses, we obtained the minimum *P*-value across variants in each locus. The gain of the multi-trait test was derived as the fraction of loci whose *P*-values were smaller than corresponding *P*-values in univariate GWAS corrected for the number of traits jointly analysed:

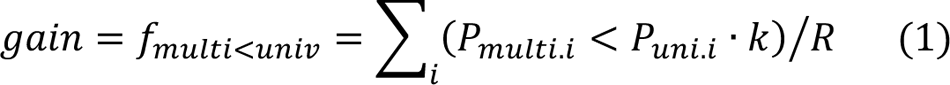

where R is the number of loci (R=1,703), *k* is the number of traits (the number of GWAS studies) jointly analysed, *P*_multi.i_ is the minimum *P*-value of the multi-trait test in region i, and *P*_*un*i*v*,i_ is the minimum *P*-value of the univariate tests across all GWAS analysed in locus i.

### Estimated and derived genetic features

We investigated the effect of both single and multi-trait features. Single GWAS features include the effective sample size (*N*_*eff*_), linear additive common variants heritability (*h*^2^_u_), polygenicity, MES, and the proportion of uncaptured linear additive common variants heritability (%*h²_u_*). Multi-trait GWAS features include the number of traits in a trait set (*k*), the average of the off-diagonal of genetic covariance and the residual covariance (Σ^’^ and Σ^’^_*r*_, respectively), condition numbers of genetic covariance matrix and residual covariance matrix (κ_)_and κ_<_, respectively) across traits, and average distance between the genetic and residual correlation matrices (Δ_+_). All parameters were aggregated to form a vector of 11 features per trait set. For the single GWAS parameters, MES, polygenicity, *N*_*eff*_, *h*^2^_GWAS_, and %*h²_u_*, we calculated mean values across each set of traits.

Polygenicity and heritability (*h*^2^_GWAS_) were estimated using MiXeR^26,27^, with the 1000 Genomes Phase3 reference panel provided along the MiXeR package containing approximately 10 million common variants^32^. Following the authors recommendation, we defined the parameter for effective sample size as *N*_*eff*_ = 1⁄(1⁄*N*_eff_ + 1⁄*N*_mult*r*mul_). For comparison purposes, we also estimated *h^2^_GWAS_* using the LDscore regression^34^, and the two metrics were consistent (Pearson ρ=0.86, **Fig. S5**). The estimated polygenicity by MiXeR showed a dependency on the GWAS sample size, with about a 10-fold increase of the polygenicity for an increase of 500,000 of the sample size (**Fig. S19**). We therefore adjusted polygenicity by taking the residuals of linear regression between log_10_ polygenicity and *N_eff_: log_10_ polygenicity(adjusted) = log_10_ polgenicity (mixer) − ケN_eff_*, where ケ was estimated by a linear regression *log_10_ ploygenicity(mixer) ∼ ケN_eff_ + ε*. We obtained ‘adjusted polygenicity’ as 10^[log_10_polynicity (adjusted)]^.We computed MES as 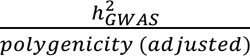

The proportion of uncaptured linear additive common variants heritability (%*h*^2^) was derived as (*h*^2^_GWAS_ − *h*^2^_GWAS-hits_)/*h*^2^_GWAS_, where *h*^2^_GWAS-hits_ denotes the heritability accounted by the univariate GWAS association loci. It was derived using the lead variants from each locus reaching genome-wide significance (*P* < 5 × 10^-8^): *h*^2^_GWAS-hits_ = ∑_i∈I_ ゲ_i_^2^, where 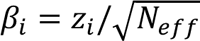. We excluded loci with lead variant whose *abs*(ゲ) > 0.19, because *h*^2^_GWAS-hits_ tended to become larger than *h*^2^_GWAS_.

The mean genetic covariance and mean residual covariance were defined as the mean of the absolute value of the upper off-diagonal elements of the genetic and residual covariance matrices (Σ_g_ and Σ_r_), i.e. σ_g_ = Σ_i,j,;i<j_σ_gij_/Σ_i,j;i<j_1, and σ_g_ = Σ_i,j,;i<j_σ_gij_/Σ_i,j;i<j_1, respectively, where σ_gij_ and σ_rij_ are ij elements of Σ_g_ and Σ_r_, and *k* is the number of traits. The condition numbers of genetic and residual covariance matrices were computed as: 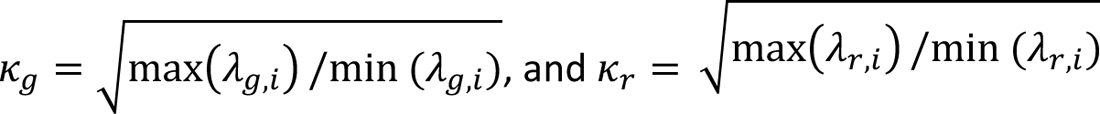 where λ_g,i_ and λ_r,i_ are the eigenvalues of the genetic and residual covariance matrices. We used *numpy.linalg.eig*^36^ for the eigen decomposition and assigned an infinite value to Σ when the minimum eigenvalue was negative or close to zero. The average distance Δ_Σ_ was defined as the mean over the absolute values of pairwise difference between the corresponding upper off-diagonal elements in genetic and residual correlation matrices Δ_Σ_ = Σ_i,j;i/j_|ρ_gij_ − ρ_rij_|/Σ_i,j,;i<j_ 1, where ρ_gij_ and ρ_rij_ indicate the ij elements in the genetic and residual correlation matrices.

### Assessment of features associated with multi-trait association gain

The contribution of features to multi-trait gain was estimated using a five-fold cross validation. For each round of cross validation, the 72 GWAS were randomly split into a training and validation data, each including 36 GWAS. Within each cross validation, we generated 1,980 unique random trait sets, containing 2 to 12 traits, for each of the training and validation data: random sampling out of the 36 traits (660 sets generated), random sampling out of traits with common SNP heritability below median (330 sets) and above median (330 sets); random sampling out of traits with MES below median (330 sets) and above median (330 sets). For this stratification of traits by the median of MES, we used polygenicity / *h*^2^_GWAS_estimated by MiXeR without adjusting for the effective sample size. For each set, we ran JASS and derived the 11 features of interest (F_i_, i = 1, …, 11) and the multi-trait gain. We used 19,793 trait sets out of the total 19,800 (=1,980 x 2 x 5) sampled for which the whole analysis process completed without error. Errors include cases where there was no association detected by both the univariate and joint tests.

Moving to the multivariate regression analysis, we selected six out of the 11 features based on collinearity analysis (as described below). The six features and the multi-trait gain were standardised into a range between 0 and 1 using *MinMaxScaler* in scikit-learn^37^. This standardisation was applied at once on the entire dataset including training and validation data across the five-fold cross validation sets.

We used the training data to estimate the joint effect δ^^^_i_ of each feature *i* from a multiple regression: gain_t*r*ain_∼ ∑_i_ δ_i_ F_t*r*ain,i_. This was conducted using the *OLS* function in statsmodels in Python^38^. We report Pearson’s correlation coefficient as a metric of predictive power.

### Collinearity and selection of features

We observed collinearity among some of the 11 features of traits across 19,266 unique trait sets. Log_10_ Σ_*r*_, log_10_ Σ_)_, log_10_ Σ^’^_*r*_, and the number of traits were highly correlated (Pearson ρ > 0.65). Likewise, mean *%h^2^_u_*, mean log_10_MES, and mean log_10_polygenicity were highly correlated (abs(ρ) > 0.8). These correlated features capture redundant characteristics of traits. Thus, we selected one out of each correlated features: the number of traits and mean *%h^2^_u_*. We chose the number of traits because it had the smallest *P*-value in a multivariate linear regression with all the features included, where inf values in condition number were replaced with their non-inf max value. We chose mean(*%h^2^_u_*) because it captures both mean(log_10_MES) (ρ=-0.90) and mean(log_10_polygenicity) (ρ=0.84). The estimation of mean(*%h^2^_u_*) is also more straightforward than mean(log_10_MES) and mean(log_10_polygenicity). After this pre-selection, we built models with the remaining six features as described in the previous section.

### Non-linear models

We considered two alternative non-linear models for prediction purposes: support vector regression (SVR), and random forest regression (RFR). SVR and RFR are regression approaches that allow for non-linear relationships. SVR’s goal is to find a hyperplane (or line, in the case of two-dimensional data) that best fits the data. It is effective at handling non-linear and complex data by using the kernel trick—mapping data with a kernel function into a higher-dimensional space where it is easier to find the best-fit hyperplane. SVR also penalises the complexity of the model and gives the flexibility in how much error is acceptable. Random forest regression performs a regression using decision trees. It generates multiple trees, fits each to a random subset of training data, and averages the predictions across trees as the final prediction. While the random sampling and averaging supposedly makes the model robust to outliers, RFR’s performance relies on a high quality of training data; the training data needs to cover a wide range as RFR does not work well for extrapolation, and RFR leads to biased predictions when the training data is sampled in a biased way^39^. In contrast, SVR is suggested to be capable of extrapolation^40^. We used the scikit-learn^37^ python implementation of RFR and of the SVR. We fitted SVR and RFR models to the gain_t*r*ain_including hyperparameters. Hyperparameters were tuned using *RandomizedSearchCV* in scikit-lean across the following range: SVR’s kernel=[linear, rbf, sigmoid, poly], C=[1,10,50,100], epsilon=[10^-3^,10^-2^,10^-1^,1], degree=[2,3,4], and RFR’s n_estimators=[5,20,50,100], max_features=[‘auto’,’sqrt’], max_depth=[12 values ranging from 2 to 100], min_samples_split=[2,5,10], min_samples_leaf=[1,2,4], bootstrap=[True, False].

### MTAG analyses

For comparison purposes, we repeated the multi-trait test and the prediction analyses with the MTAG approach^4^. MTAG uses a weighted sum of *Z*-score (**Supplementary note 1**). Trait weights are derived using the generalised method of moments, as (ゲ^y^_I_ゲ^yR^− Ω − Σ_i_) = 0, where Ω is the genetic covariance matrix to be estimated for the weights, and Σ_i_ is the genetic covariance matrix estimated using the LD score regressions by Bulik-Sullivan et al^34^. The model assumes that the genetic covariance matrix is homogenous across variants. We used MTAG with its default setting, which considered only complete cases. We ran MTAG for the same trait sets used for the analysis with JASS (all data used in the five-fold cross validation). MTAG outputs *P*-values for each trait in each trait set, whereas JASS gives a single *P*-value for a trait set. To account for the number of tests run by MTAG for one set, we obtained minimum *P*-values by MTAG across traits and variants in each locus and multiplied the minimum *P*-values by the number of traits in the set. For the comparison of the association gain between MTAG and JASS, we used both the minimum *P*-values with and without multiple test correction (**Fig. 5**, and **Fig. S14**).

### Comparison of strategies for trait selection

To compare the performances of trait selection strategies, we classified the 19,266 unique sets into clinically homogenous, low heterogeneity, high heterogeneity, or high predicted gain according to our predictive model. To assess clinical homogeneity, we first classified the 72 traits into clinical groups using the broadest categories in the MeSH Tree Structures^41^. The grouping was further refined based on clinician’s insights (**Table S1**). We labeled each set of traits as ‘homogeneous’ if all the traits are in the same clinical group, ‘low heterogeneity’ if trait belonged to two to four clinical groups, ‘high heterogeneity’ if traits spanned five or more clinical groups. For the ‘data-driven’ method, we selected 100 sets of traits by CV fold that had the largest gains predicted by the linear model with aggregated coefficients across CV folds (**Table 1**). We used the Welch’s *t*-test to evaluate the differences in gain and the number of new associated loci. We used a two-sided Welch’s *t*-tests to determine whether the data driven method achieves a greater association gain than other methods, and whether jointly analysing clinically heterogenous traits achieves a greater association gain than jointly analysing clinically homogenous traits. For this test, we used a pair of training data and validation data that are mutually exclusive to ensure independence.

### Evaluation of the relevance of new association

To evaluate the relevance of new associations detected by JASS (i.e. if most of them were true positive), we attempted to predict loci discovered in a recent large meta-analysis on BMI (sample size of 683,365 on average across ∼2.3 million variants^12^) from the results of multi-trait GWAS applied on 1,776 trait sets containing a smaller study of BMI (sample size of 339,224, **Table S1**). First, we compared loci detected by JASS (after a Bonferroni correction to account for the number of sets) and in the larger GWAS using the standard genome wide significance threshold of 5 × 10^-8^. Second, we fitted a logistic regression to predict associated loci in the larger GWAS by combining JASS *P*-value and the number of sets where the loci was considered associated with JASS. We use odd number chromosomes to fit the logistic regression and evaluated its performances on even number chromosomes.

