## Supplemental information for "Trait selection strategy in multi-trait GWAS: Boosting SNPs discoverability"

### Supplementary Material

#### Supplementary Notes

##### *1. Standard Multi-trait approaches: the Omnibus test and weighted sum test*

Amongst the many existing multi-trait methods, some posit a normal distribution under the null hypothesis whose parameter depends on sample overlap between traits. As a variance testing framework, two methods have been often used, namely a $k$ degree of freedom omnibus test, where $k$ equals the number of traits analysed jointly [1–3] and 1 degree of freedom weighted sum of single trait statistics, where the weights are defined based on a priori of the direction and importance of each trait[4–6]. The omnibus test has been recently suggested to perform better than the weighted sum of *Z*-scores[7].

The high performance of the omnibus test can be explained as it uses the combination of weighted *Z*-scores; while a version of the weighted sum of *Z*-scores (implemented in JASS) uses one of the eigenvectors of the estimated covariance matrix as the weight, the omnibus test is equivalent to the sum of this weighted *Z*-scores across all eigenvectors[7]. Moreover, while the weighted sum test requires weights that require additional assumptions, such as a homogeneous genetic across variants[6], the omnibus test does not require such assumptions that could be met only under limited cases (**Fig. S3**).

While some of these multi-trait GWAS use different ways to estimate the covariance matrix under the null, there is now a well-established method using LDscore regression[8], which has been incorporated by more recent multi-trait GWASs[1,5,6,9]. Taking advantage of these studies, here we use the omnibus test with the null covariances estimated by the LDscore regression as the main approach.

#### *2. Influence of genetic and residual covariances on multi-trait (omnibus) test performance*

To assess the impact of traits on the statistical power of multi-trait GWAS, one of the metrics we used is the genetic-residual distance as a measure to characterise the orientation difference between the residual covariance and genetic covariance. The orientation difference can be associated with the statistical power gain in the omnibus test because the null in the omnibus test is in k-dimensional space (k=the number of traits), where not only the magnitude of *Z*-scores but also its angle determines whether the corresponding variants are significant (**Fig. S3**). This suggests that a larger orientation difference (e.g. larger genetic-residual distance) is associated with increased statistical power. We also measured two properties of each covariance matrix individually using mean and condition number.

#### *3. Stratification in trait sets sampling*

In the generation of trait sets, we used random sampling out of all 72 traits (split into two datasets for training and validation) and sampling with stratification of traits (**Material and methods**). Random sampling will result in trait sets whose mean of genetic features are similar across trait sets. Indeed, the variance of the mean is smaller than the variance of a sampled random variable (the variance of the empirical mean is $var(\bar{X}) = var(X)/N$ where *X* denote a random variable and *N* the number of samples). Such a narrow range for genetic feature could impair regression analysis.

To improve regression analysis with a wider range of mean genetic features as explanatory variables, we sampled trait sets with three methods: (i) random sampling, (ii) stratified sampling with mean effect size and (iii) stratified sampling with heritability. The right panel represent the distribution of a random variable sampled from a uniform distribution (red), the distribution of the mean of the variable across sets generated with random sampling (yellow), and the distribution of the mean of the variable across sets generated with the stratification (blue). Note that the largest trait sets we generated contain *k*=12 traits. This is because the data split for training and validation and stratified by median of a feature becomes ~1/4 of the original size (1/4 x 72 GWAS = 18), requiring $k < 18$. $k$=12 allowed us to generate a number of distinct trait sets as large as 1,980 sets.

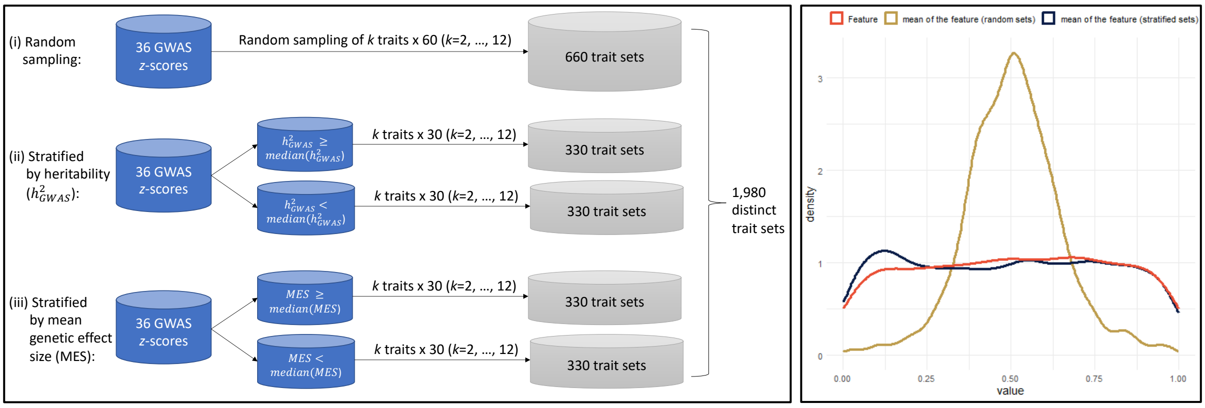


#### 4. *A hypothetical example of a negative contribution of $\Delta_\Sigma$ to the multi-trait (omnibus) gain.*

We visualise a hypothetical example where an increase in $\Delta_\Sigma$ leads to a decrease in the omnibus gain, the suggested trend from our analysis with the 72 GWAS.

Our analysis revealed two trends about genetic covariance and residual covariance: (i) Small $\Delta_\Sigma$ (**Fig. 2**) implies the distributions of genetic effect size distribution (blue ellipse) and the omnibus null distribution (green ellipse) are oriented roughly in the same direction; (ii) on average, mean genetic correlation was greater than mean residual correlation across 19,266 unique trait sets (Panel A), suggesting the blue ellipse is narrower than the green one below.

Panel B visualizes hypothetical distributions of the effect size and the omnibus null with the two trends satisfied. Asterisks indicate the areas corresponding to the gain by the omnibus test relative to univariate test, whose thresholds are shown as the square; the part of the blue ellipse that is inside the square and outside the green ellipse indicates the gain.

The multivariate model we built suggested a negative impact of $\log_{10}\Delta_\Sigma$ on multi-trait gain while controlling for other features including $\log_{10}\bar{\Sigma}_g$ and mean($h^2_{GWAS}$). In other word, keeping the blue ellipse fixed. A change that corresponds to an increase in $\log_{10}\Delta_\Sigma$, while the distribution of genetic effect size being fixed, is changing the correlation in the omnibus null distribution such that the correlation becomes more different from the correlation in genetic effect size distribution than before changing. Such a change corresponds to reducing the correlation in the omnibus null from panel B to C, while everything else is kept the same (note that the areas of the omnibus null distributions in panel B and C are also constant). The area marked by asterisks in panel B reduced in panel C.

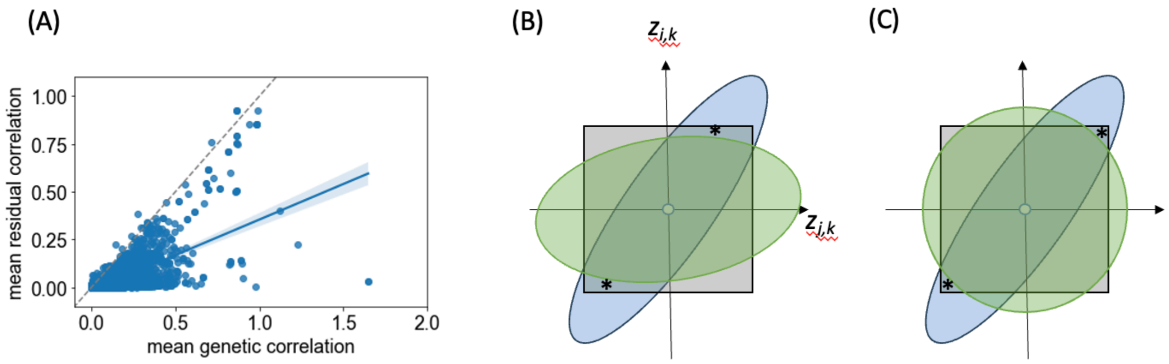

***Figure S1. Genetic correlation matrix across 72 traits.***

Pairwise genetic correlation derived using the LDscore regression between each of the 72 traits. Traits were reordered using a hierarchical clustering (complete linkage, distance= 1- ρ). (q-value annotation: *** < 10-3, ** <10-2,*< 5x10-2, .< 10-1)

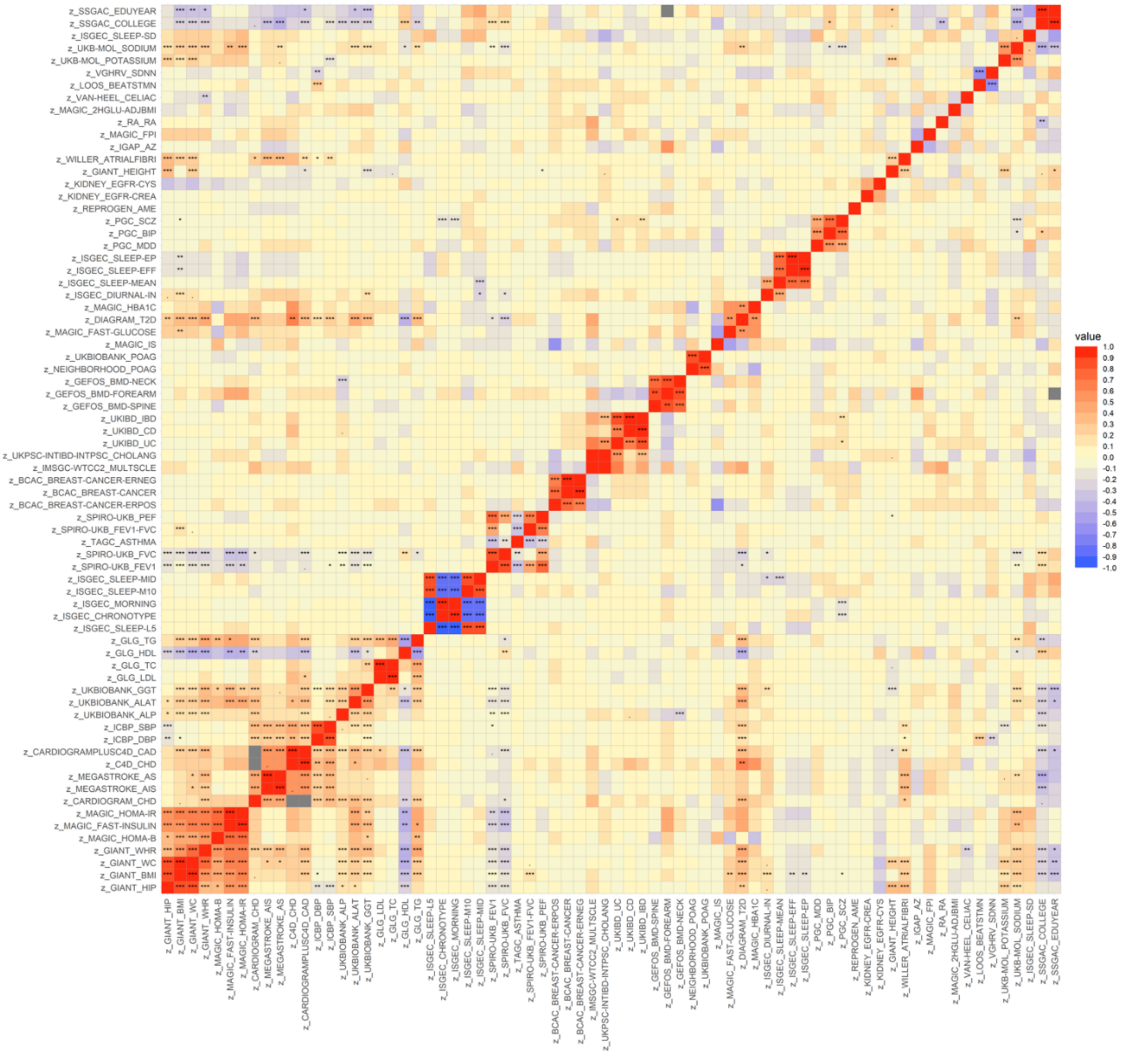

***Figure S2. Residual covariance matrix across 72 traits***

Residual covariance derived using the LDscore regression between each of the 72 traits.

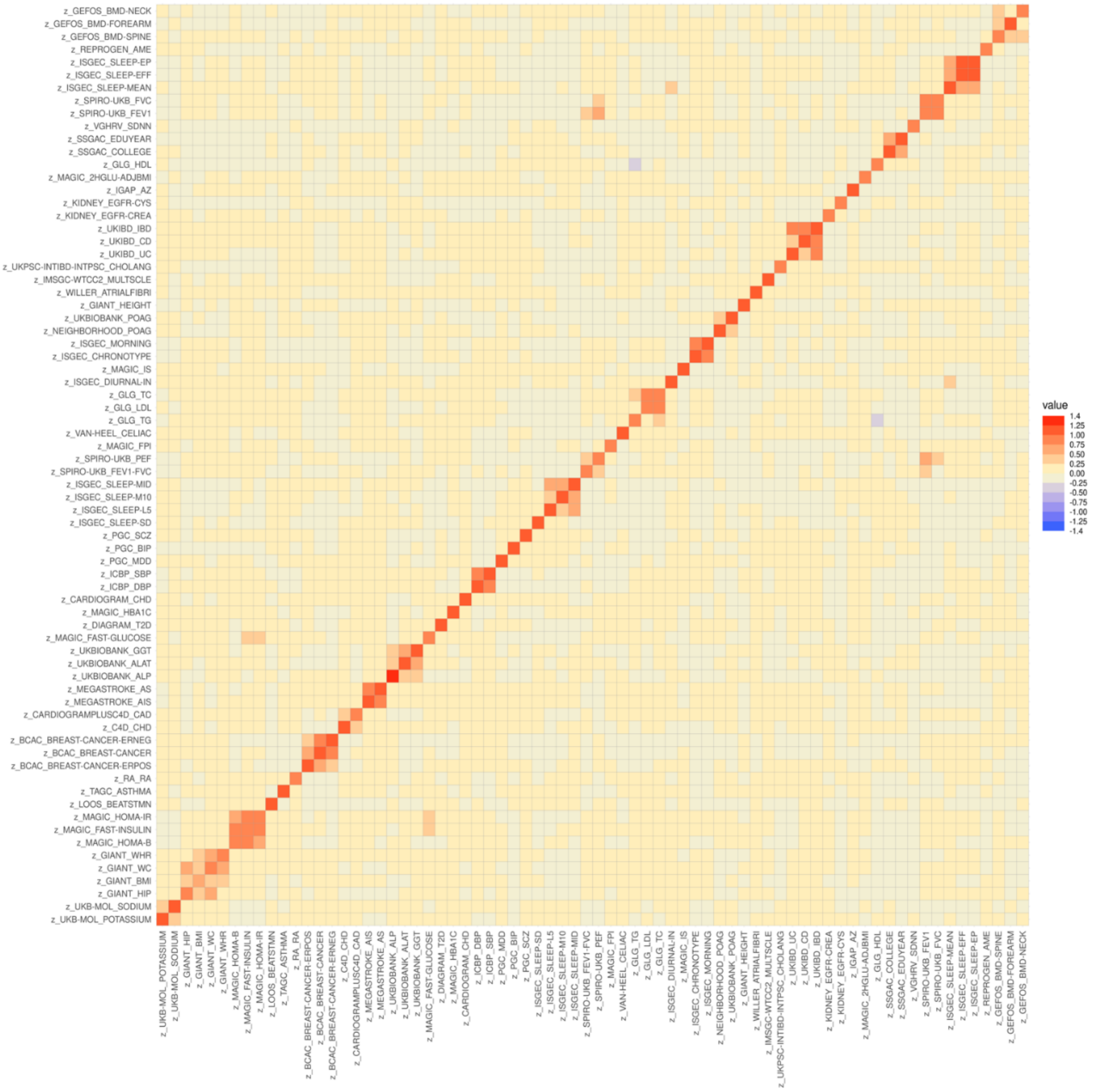

***Figure S3. Schematic of the omnibus test and its gain over the univariate test on several bivariate examples.***

The blue dots and shades represent *Z*-scores of SNPs in k=2-dimensional space (k: the number of traits jointly analysed). The black square indicates the significance threshold of the standard univariate GWAS. The green ellipse depicts the significance threshold of the omnibus test. $\hat{\beta}_{i,k}$: coefficient from a GWAS for $SNP_i$ for trait $_k$, $\hat{\sigma}_{i,k}$: standard error of $\hat{\beta}_{i,k}$.

(A-C) The three panels visualise cases where the residual covariance matrix ($\Sigma_r$) captures (A) positive correlation, (B) no correlation, and (C) negative correlation between traits. SNPs that are inside the square and outside the green ellipse are significant for the omnibus test but not for the univariate test (e.g. A). In contrast, SNPs that are inside the green ellipse but outside the black square are significant for univariate tests but not significant for the omnibus test (e.g. C).

(D-F) Instead of a single SNP, the distribution of *Z*-scores from GWAS are shown with different genetic covariance ($\Sigma_g$). Given the positive covariance in the null distribution, the omnibus test has a larger gain when genetic covariance is 0 (e.g. D) or negative and misaligned with residual covariance (e.g. E), whereas the omnibus test has a smaller gain when genetic covariance is strongly positive and aligned with residual covariance (e.g. F). Note that the omnibus test can yield a large gain relative to univariate tests even when there is no genetic correlation between traits as shown in (D).

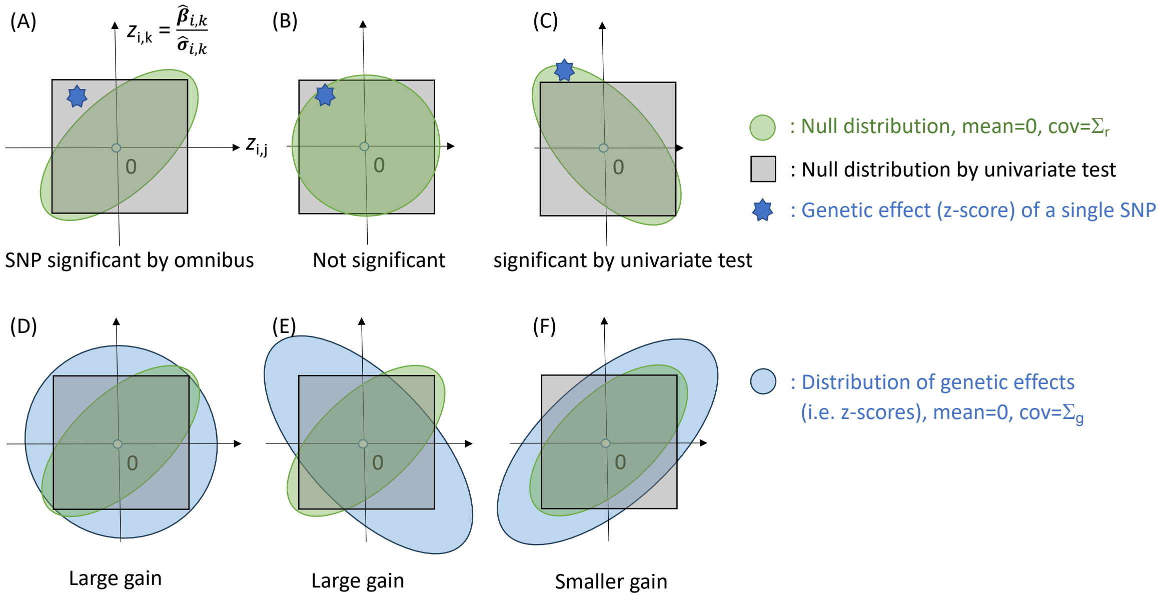

(A)
$z_{i,k} = \frac{\hat{\beta}_{i,k}}{\hat{\sigma}_{i,k}}$
$z_{i,j}$
0
SNP significant by omnibus
(B)
0
Not significant
(C)
0
significant by univariate test
: Null distribution, mean=0, cov=$\Sigma_r$
: Null distribution by univariate test
: Genetic effect (z-score) of a single SNP
(D)
0
Large gain
(E)
0
Large gain
(F)
0
Smaller gain
: Distribution of genetic effects
(i.e. z-scores), mean=0, cov=$\Sigma_g$

***Figure S4. Distributions of five features with and without log10 transformations.***

(A) Violin plot of the distribution of feature $\bar{\Sigma}_g, \bar{\Sigma}_r, \Delta_{\Sigma}, \kappa_g, \kappa_r$. (B) Violin plot of the distribution of the same features log-transformed.

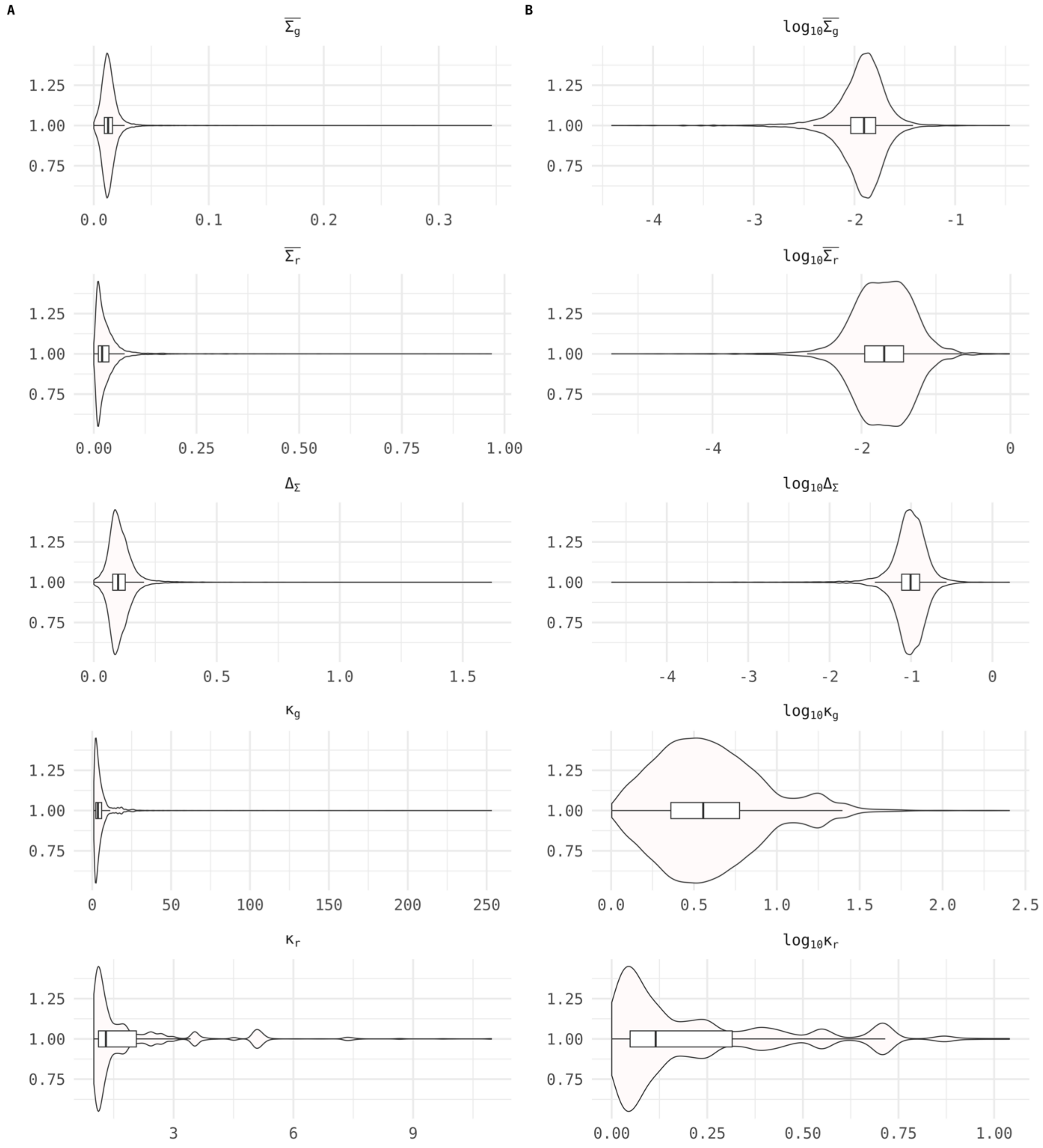

***Figure S5. Comparison of heritability between MiXeR and LD score regression***

For each of the 72 traits we estimated the heritability on the observed scale using both MiXeR and LD score regression.

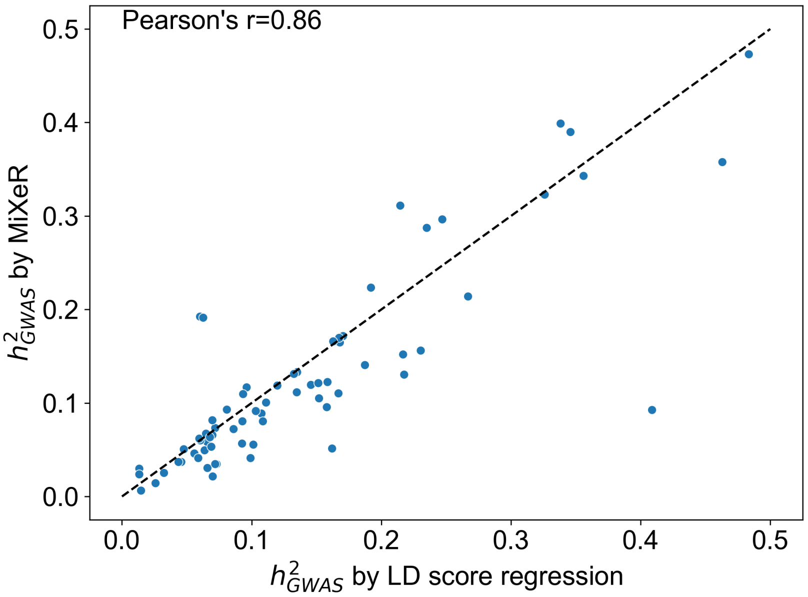


***Figure S6. Relationship between the fraction of undetected heritability ($\%h_u^2$), linear additive heritability of common variants ($h^2_{GWAS}$), polygenicity and mean genetic effect size.***

Scatter plot of the fraction of undetected heritability ($\%h_u^2$) with respect to heritability of common variants ($h^2_{GWAS}$). Dot colors represents polygenicity in A), and mean genetic effect size in B)

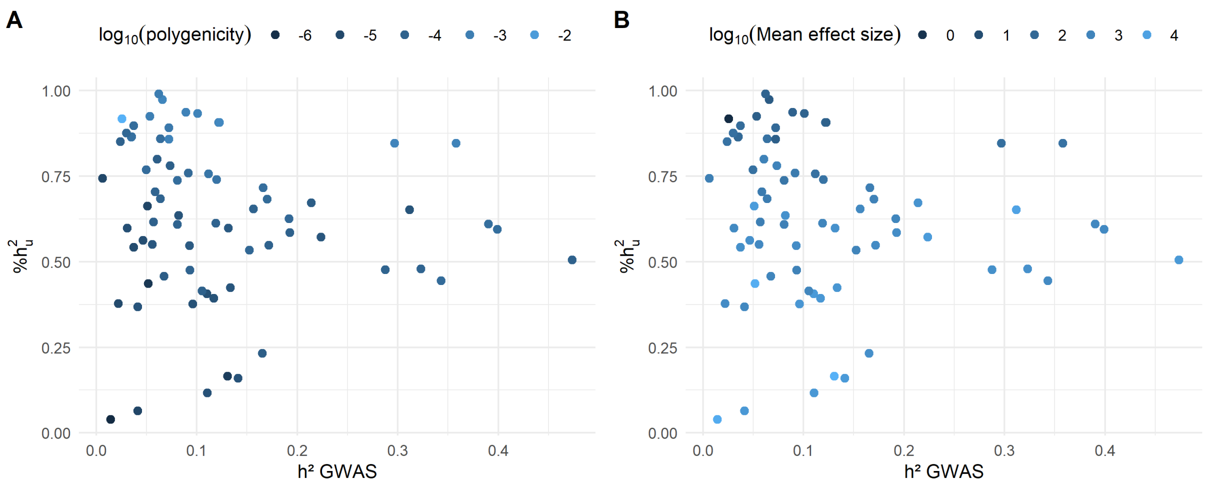

***Figure S7. Relation between the number of loci detected by the multi-trait test, the gain, and the number of loci previously detected by univariate tests.***

A) Percentage of significant associations that were detected by JASS with respect to the association gain. B) Number of loci detected by the multi-trait test with respect to the number of loci previously associated with the trait (significant for one of the univariate tests). Dot colours represent the association gain. C) Histogram of the number of new associations detected by JASS when applied on variants with on observed genetic effect for all traits. D) Histogram of the number of new associations detected by JASS when applied all variants including those with missing genetic effect for a subset of traits.

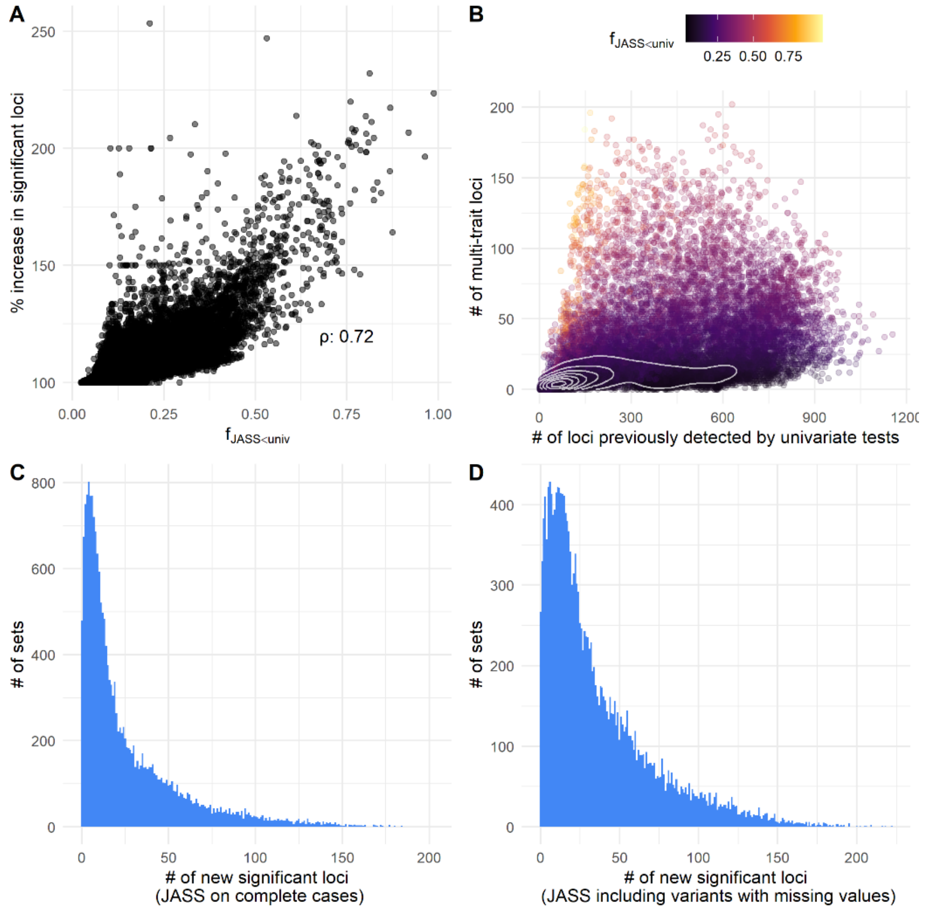

***Figure S8. The relationship between the number of new associations and gain of JASS and the condition number of the residual covariance matrix.***

Each dot represents a trait set on which JASS performed a multi-trait GWAS (Omnibus test). Note that the dots on the right most side of the figure (log10 condition number > 1.2) are cases when the minimum eigenvalue of the covariance matrix was negative (and hence flagged as a singular matrix), for which assigned Inf condition number. Here, we replaced Infs with 2 x non-Inf maximum condition number for evaluation purpose.

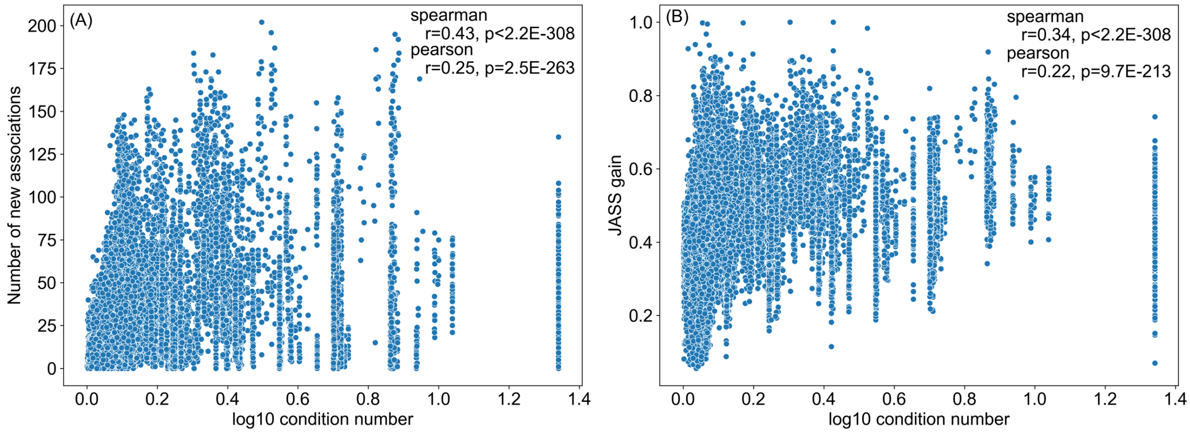

***Figure S9. Overview of the five-fold cross validation processes.***

*Z*-scores from 72 GWAS are denoted as $z_j$ $(j = 1, \cdots, 72)$. For each of the trait set generated, we obtained six genetic features. Here, genetic features obtained for trait sets in training and validation data are denoted as $F_{train,i}$ and $F_{val,i}$ $(i = 1, \cdots,\ 6)$, respectively, and these six features are $mean(\%h_u^2)$, *#traits*, $mean(\log_{10} \Delta_\Sigma)$, $mean(\log_{10} \bar{\Sigma}_g)$, $mean(N_{eff})$, and $mean(h_{GWAS}^2)$. $\hat{\delta}_i$ indicate the estimated joint effect from the multiple regression $f_{JASS<univ,train} \sim \sum_i \delta_i F_{train,i}$. We measured Pearson's correlation coefficient ρ to evaluate the performance of the regression model.

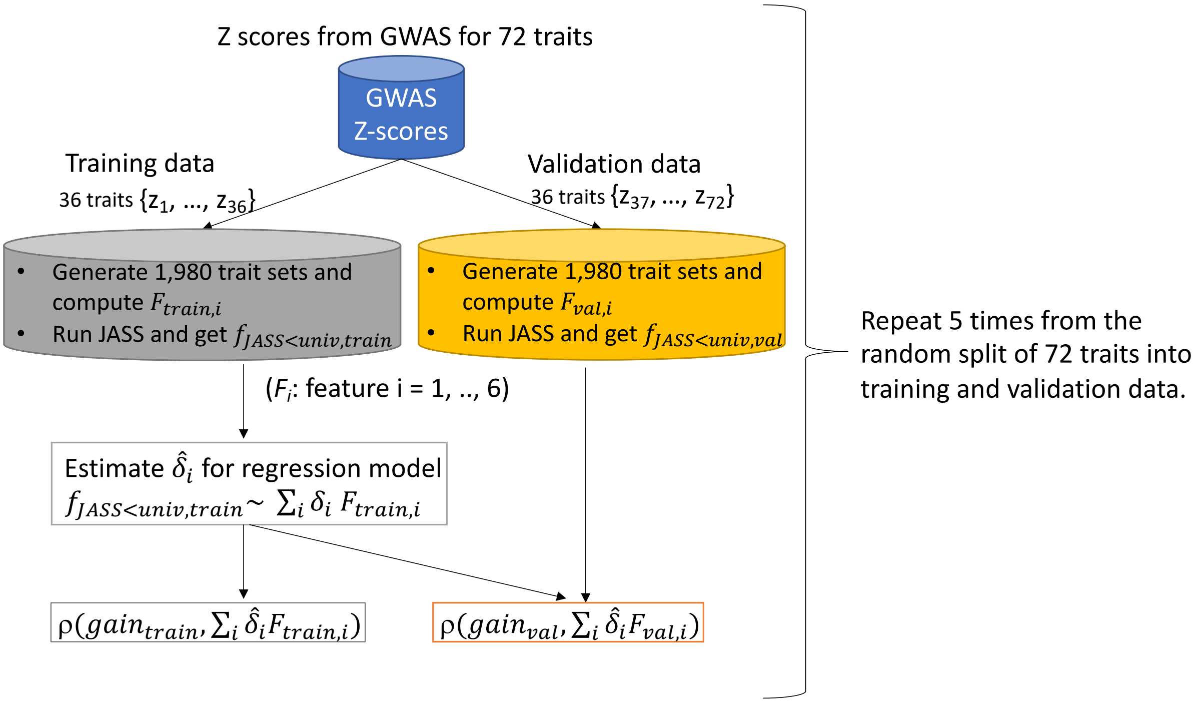


***Figure S10. Observed JASS gain vs predicted gain for validation data.***

*The predicted gain was obtained from the linear model with the pre-selected features.* Vertical and horizontal grey lines show the mean values of the predicted and observed gains, respectively.

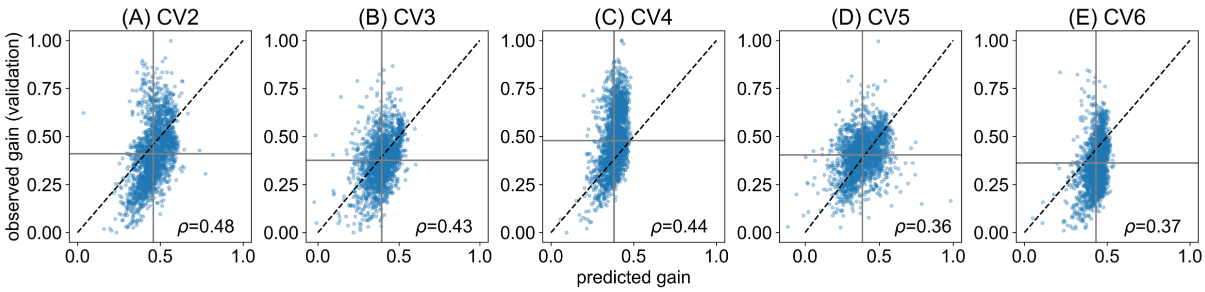

***Figure S11. Regression analysis summary with mean polygenicity and mean MES instead of*** ***%$h^2_u$.***

Panels (A-C) show the performance of the model with mean polygenicity, and (D-F) show the performance with mean MES, replacing mean $\%h_u^2$ in **Figure 4**. (A, D) Boxplots of the prediction power across the five-fold cross validations (CV) of the multivariate linear regression model measured as the Pearson's correlation coefficient between the predicted and observed gain. The performance of each CV is represented as a coloured dot. Orange dashed line: median correlation coefficient between the predicted and observed gain in the validation data. (B, C, E, F) The boxplots show the coefficients and -$\log_{10}$(*P*-values) of the six features in the regression model across five-fold cross validations using each corresponding training data. Red dashed line: Bonferroni corrected nominal significance threshold.

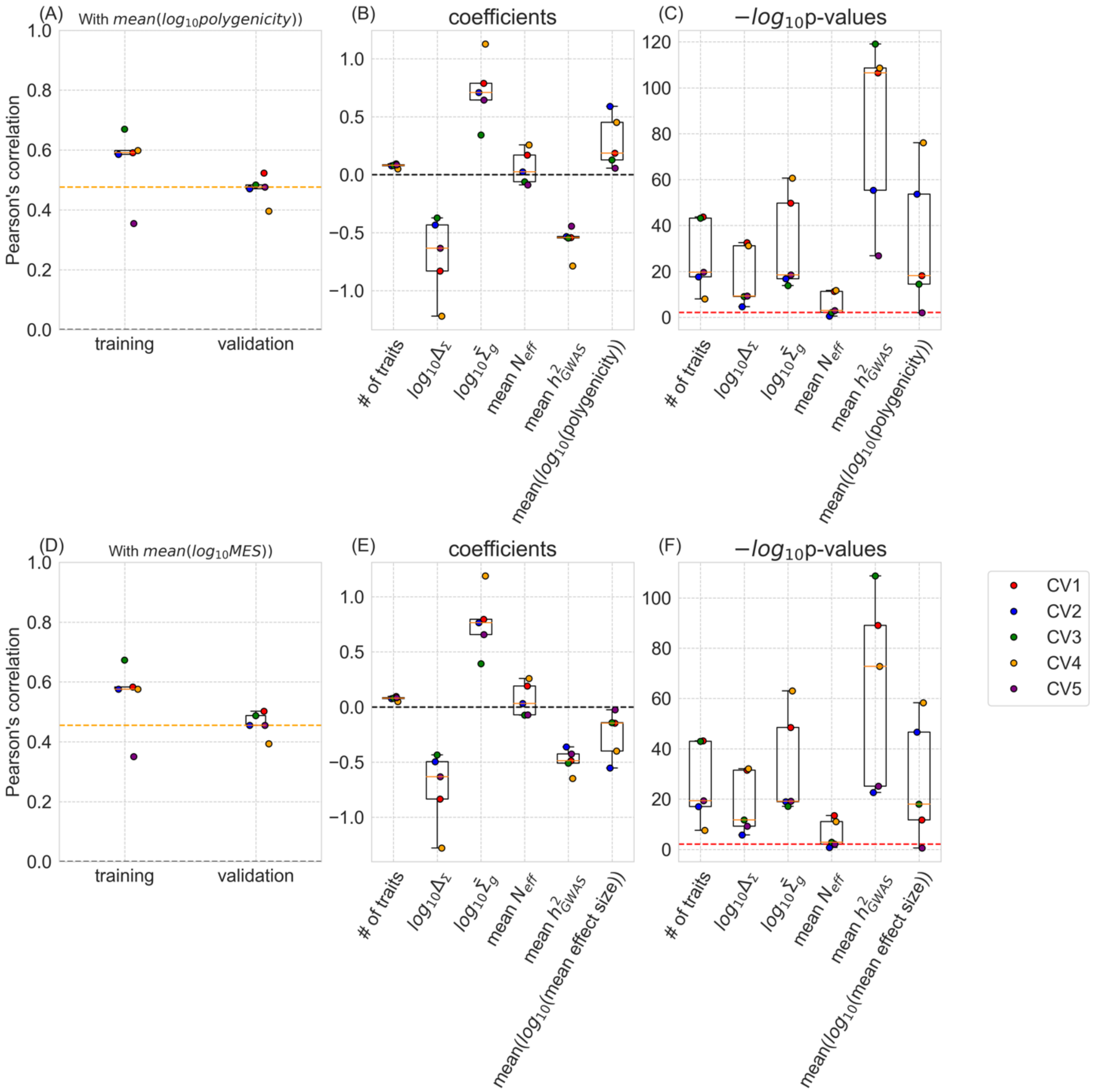
(A)
With $mean(log_{10}polygenicity)$)
Pearson's correlation
1.0
0.8
0.6
0.4
0.2
0.0
training
validation
(B)
coefficients
1.0
0.5
0.0
−0.5
−1.0
(C)
$-log_{10}$p-values
120
100
80
60
40
20
0
### of traits
$log_{10}\Delta_{\Sigma}$
$log_{10}\hat{\Sigma}_g$
mean $N_{eff}$
mean $h^2_{GWAS}$
$mean(log_{10}(polygenicity))$
(D)
With $mean(log_{10}MES)$)
Pearson's correlation
training
validation
(E)
coefficients
(F)
$-log_{10}$p-values
$mean(log_{10}(mean\ effect\ size))$
CV1
CV2
CV3
CV4
CV5

***Figure S12. Prediction power of non-linear models.***

We measured the prediction power of the multivariate non-linear regression model, namely support vector regression (SVR) and random forest regression (RFR), as the Pearson's correlation coefficient. Results from five-fold cross validations are shown as box plots. Orange dashed line: median correlation coefficient between the predicted and observed gain in the validation data.

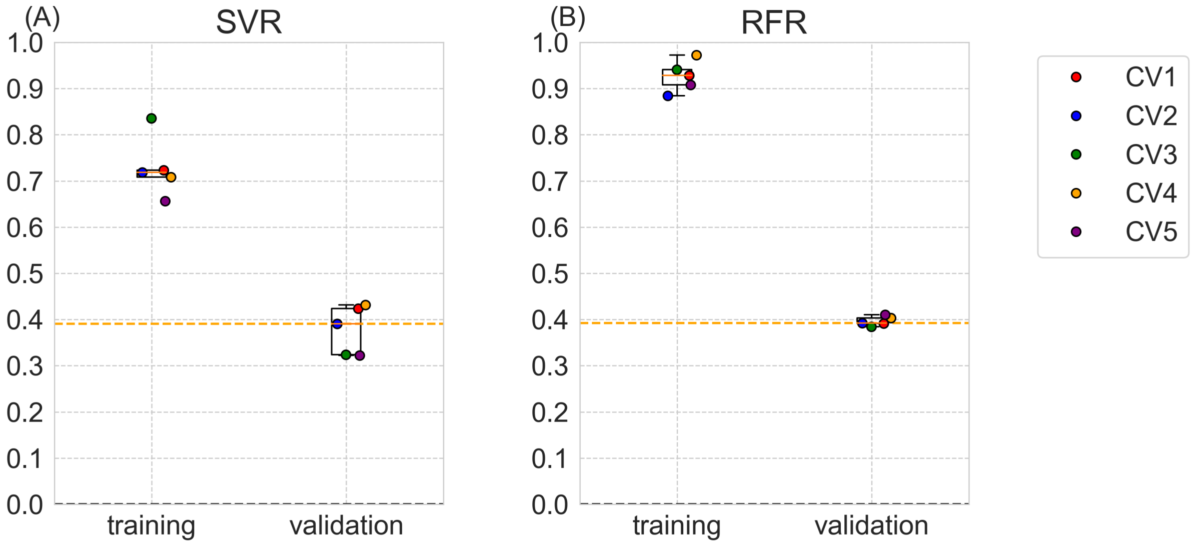

***Figure S13. Regression analysis summary for MTAG association gain.***

(A) Boxplots of the prediction power across the five-fold cross validations (CV) of the multivariate linear regression model measured as the Pearson's correlation coefficient between the predicted and observed MTAG gain. The performance of each CV is represented as a coloured dot. Orange dashed line: median correlation coefficient between the predicted and observed gain in the validation data. (B, C) The boxplots show the coefficients and -$\log_{10}$($P$-values) of the six features in the regression model across five-fold cross validations using each corresponding training data. Red dashed line: Bonferroni corrected nominal significance threshold.

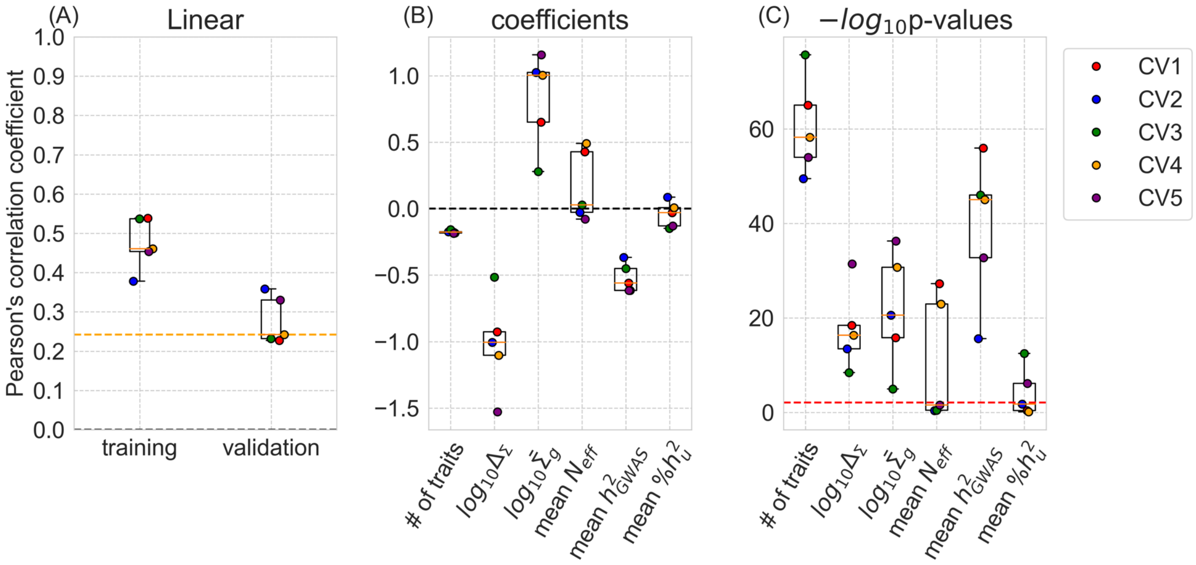

***Figure S14. Comparison of MTAG (uncorrected for multiple testing) with JASS.***

A) Numbers of new associations found by JASS with respect to the number of new associations found by MTAG (uncorrected for multiple testing) across all the trait sets. Each dot represents a set of traits. Dot color represents the number of traits in the set. B) Fraction of sets where the number of new associations detected by JASS was superior to the number of associations detected by MTAG stratified by the number of traits in the set.

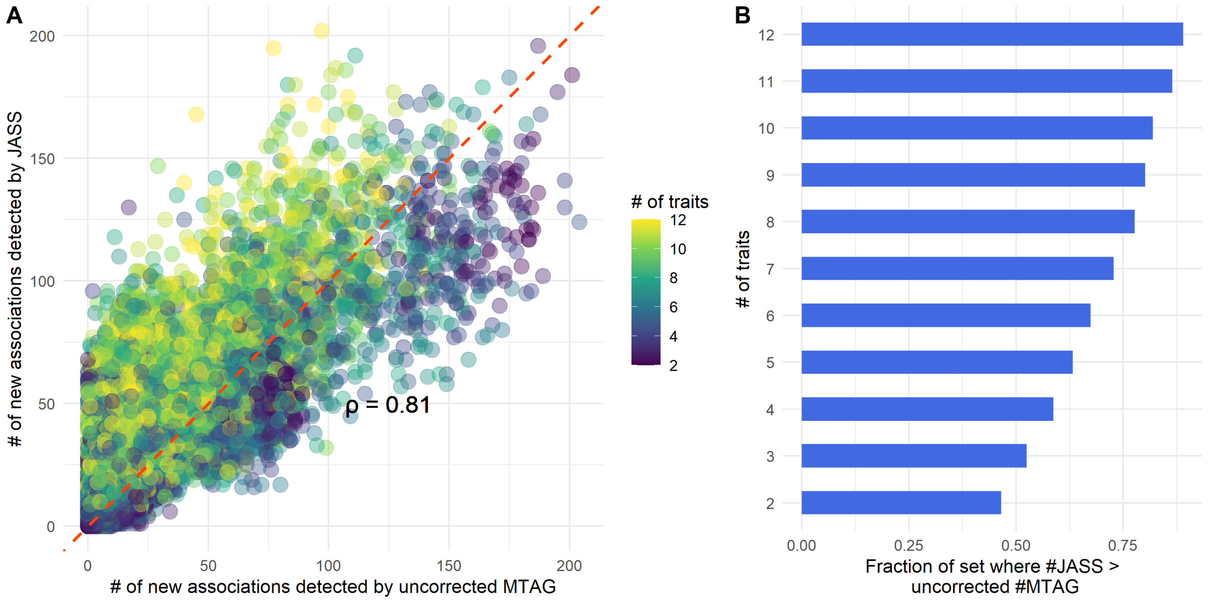

***Figure S15. Barplot of the Fold enrichment of traits in the top 10% of sets (in terms of new associations detected by JASS) against the 10% of bottom sets.*** *Traits are referred by their acronyms, refer to Table S1 for their full names.*

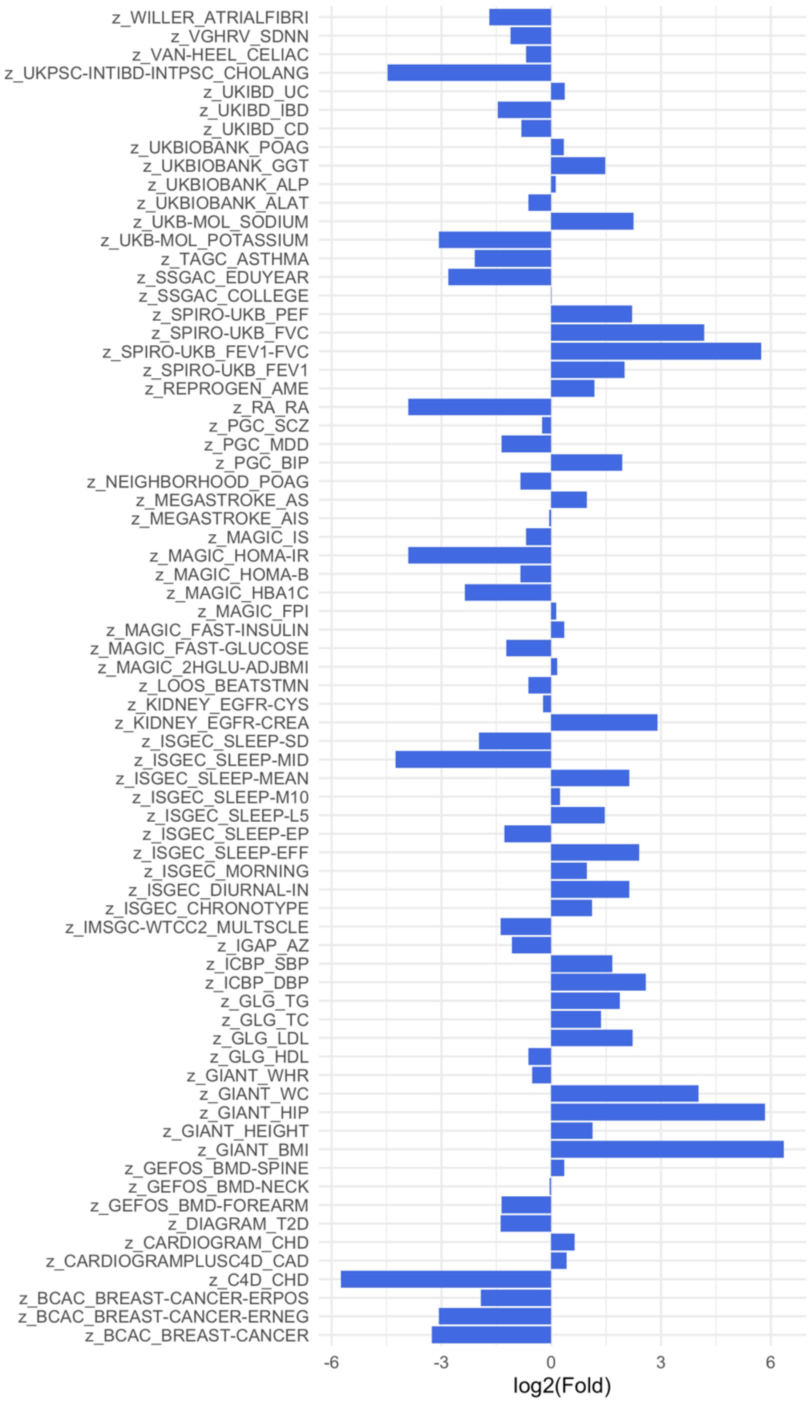

***Figure S16. Receiver Operating Characteristic (ROC) curve for the prediction of new associations of a larger GWAS study on BMI* (sample size= 683,365) *based on multi-trait associations detected using a smaller BMI study* (sample size= 339,224).**

ROC curves were derived using three continuous makers for putative BMI loci derived from the 1,776 multi-trait GWAS tests on sets containing BMI: i) the number of sets where the loci was associated, ii) the minimum *P*-value observed across sets (-log10(minimum *P*-value))., iii) probability of being detected in the larger GWAS based on a logistic regression combining the minimum JASS *P*-value across set, and number of significant sets. Curves colors correspond to marker.

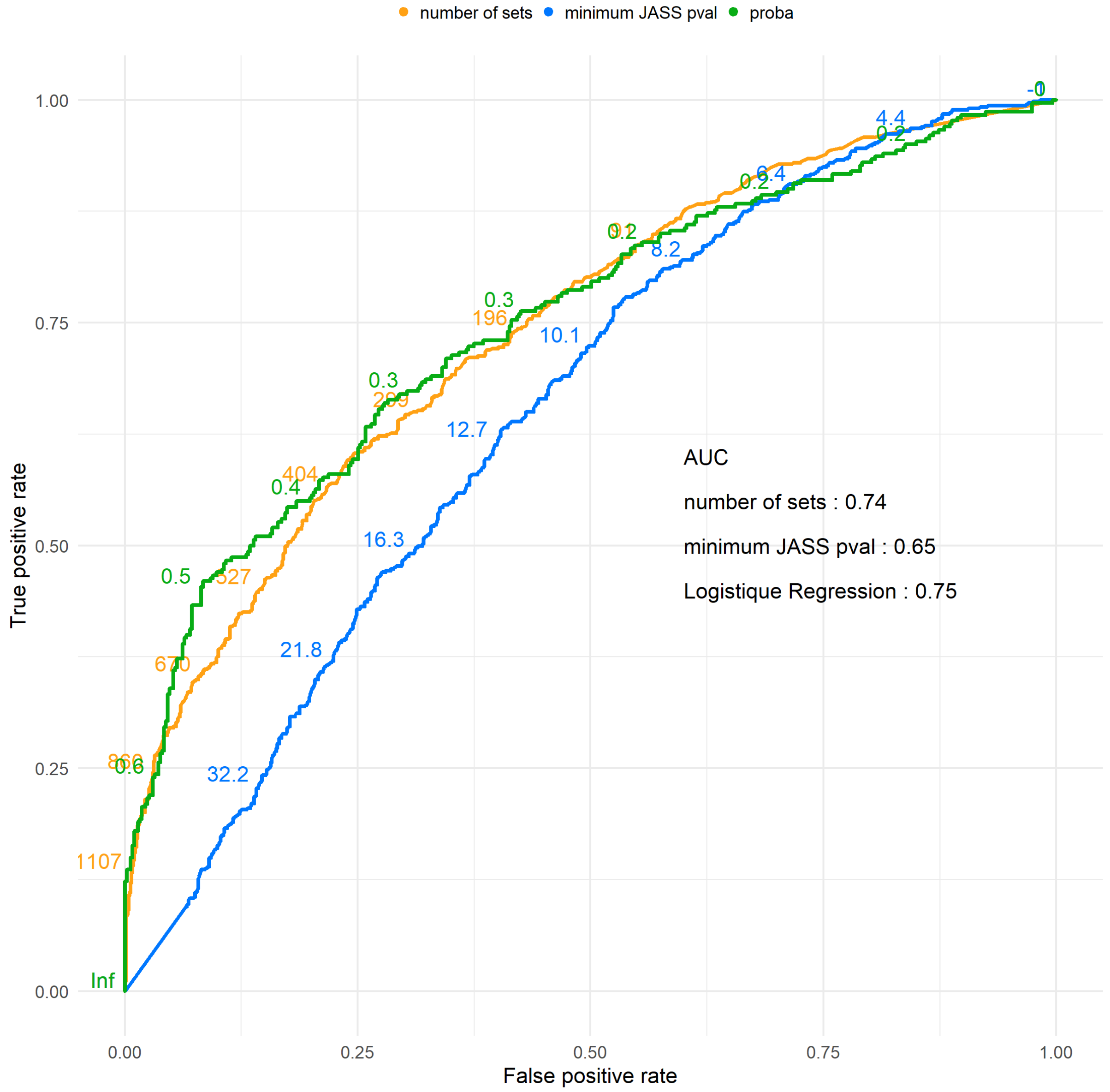

**Figure S17. Comparison between clinical and data-driven trait sampling methods when applying MTAG.**

(A, B) Distribution of the gain and the number of new association loci for trait sets selected by four trait selection strategies from the validation data. *P*-values are from the two-sided Welch's *t*-test. Differences in mean values in each pair compared in the test (right – left categories in the order shown on the x-axis) are also shown. Note we used a pair of a training data and a validation data to ensure independence for the test. The numbers under the labels on the x-axis indicate the number of trait sets from each strategy. The observed MTAG gain and the number of new association loci are shown on the y-axis. (C, D) Comparison between MTAG and JASS in terms of gain and the number of new association loci for 'homogenous' trait sets. (E, F) Comparison between MTAG and JASS in terms of gain and the number of new association loci for trait sets of the 'homogenous' category, visualised per clinical grouping.

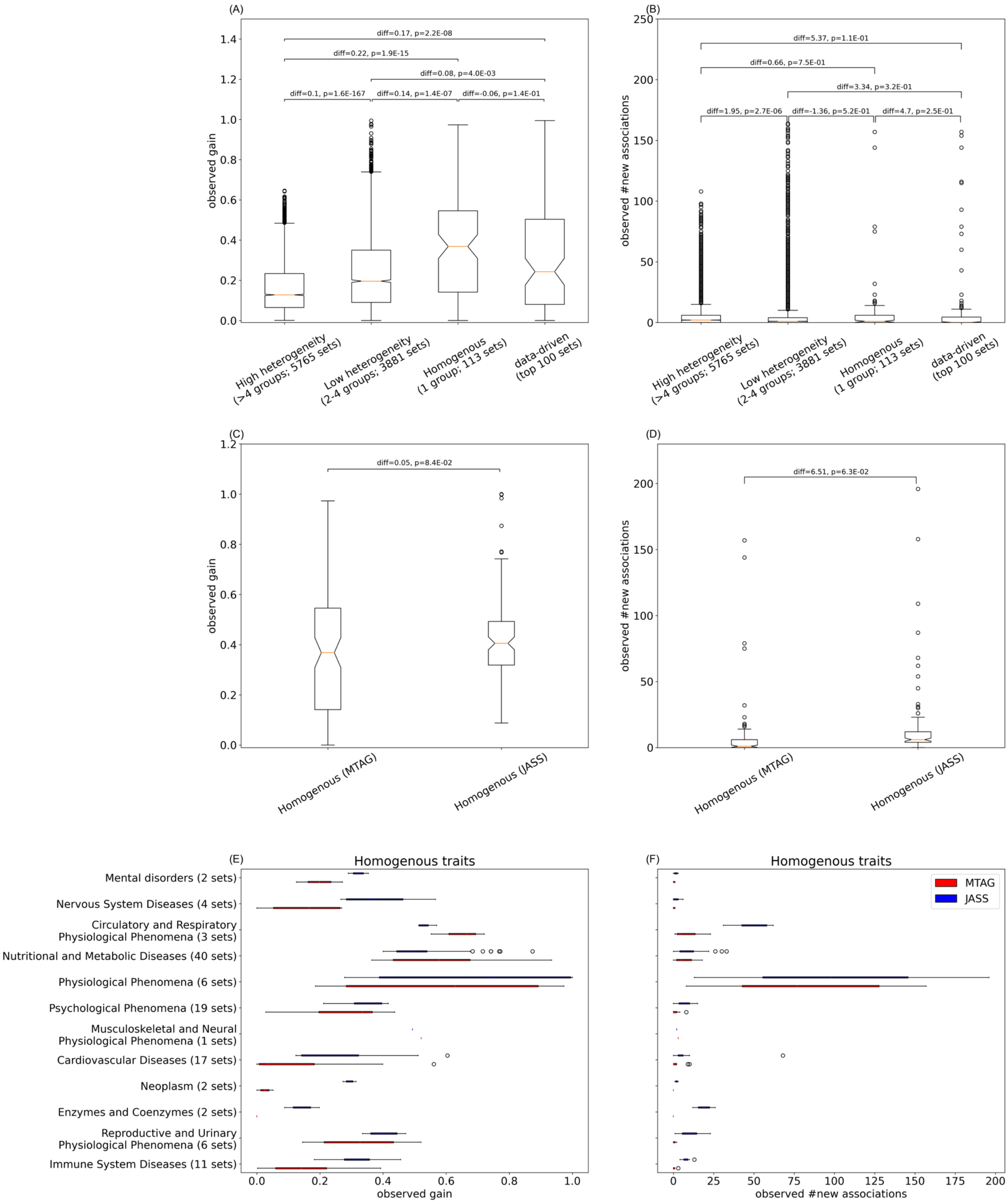
(A)
diff=0.17, p=2.2E-08
diff=0.22, p=1.9E-15
diff=0.08, p=4.0E-03
diff=0.1, p=1.6E-167
diff=0.14, p=1.4E-07
diff=-0.06, p=1.4E-01
observed gain
High heterogeneity (>4 groups; 5765 sets)
Low heterogeneity (2-4 groups; 3881 sets)
Homogenous (1 group; 113 sets)
data-driven (top 100 sets)
(B)
diff=5.37, p=1.1E-01
diff=0.66, p=7.5E-01
diff=3.34, p=3.2E-01
diff=1.95, p=2.7E-06
diff=-1.36, p=5.2E-01
diff=4.7, p=2.5E-01
observed #new associations
High heterogeneity (>4 groups; 5765 sets)
Low heterogeneity (2-4 groups; 3881 sets)
Homogenous (1 group; 113 sets)
data-driven (top 100 sets)
(C)
diff=0.05, p=8.4E-02
observed gain
Homogenous (MTAG)
Homogenous (JASS)
(D)
diff=6.51, p=6.3E-02
observed #new associations
Homogenous (MTAG)
Homogenous (JASS)
(E)
Homogenous traits
Mental disorders (2 sets)
Nervous System Diseases (4 sets)
Circulatory and Respiratory Physiological Phenomena (3 sets)
Nutritional and Metabolic Diseases (40 sets)
Physiological Phenomena (6 sets)
Psychological Phenomena (19 sets)
Musculoskeletal and Neural Physiological Phenomena (1 sets)
Cardiovascular Diseases (17 sets)
Neoplasm (2 sets)
Enzymes and Coenzymes (2 sets)
Reproductive and Urinary Physiological Phenomena (6 sets)
Immune System Diseases (11 sets)
observed gain
(F)
Homogenous traits
MTAG
JASS
observed #new associations

***Figure S18. Suggested approaches depending on the trait selection strategy.***

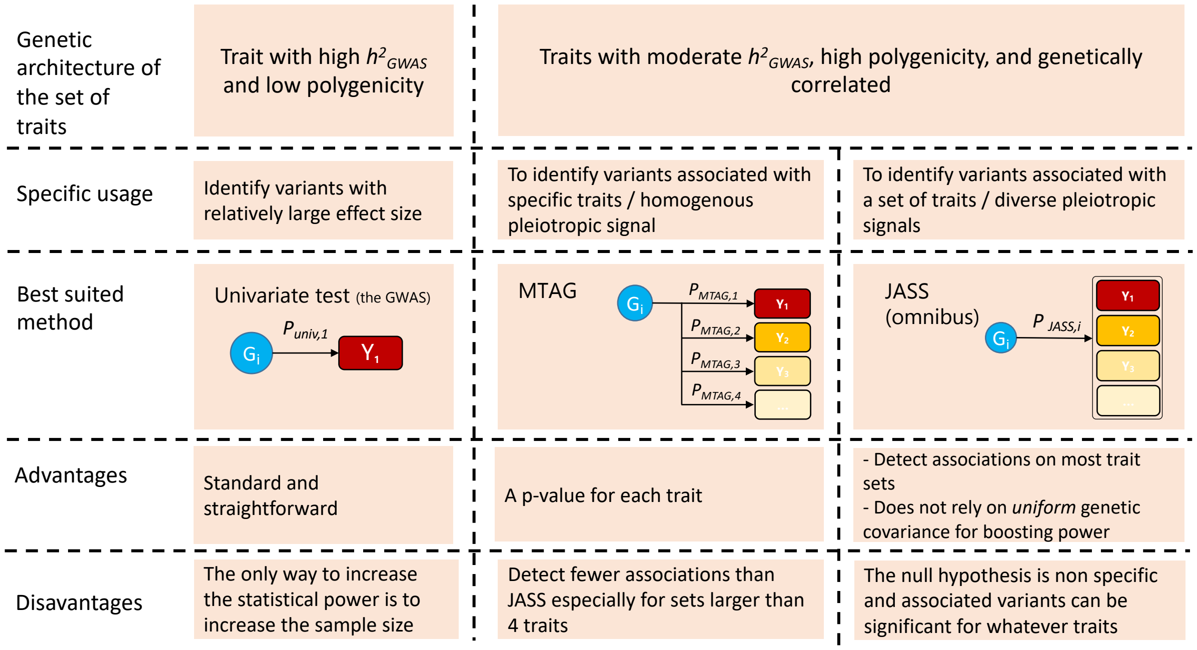


***Figure S19. Dependency of the polygenicity estimate with sample size.***

For each of the 72 traits we plotted the polygenicity estimated by MiXeR as a function of the reported effective sample size. Regression line and 95% confidence interval are shown.

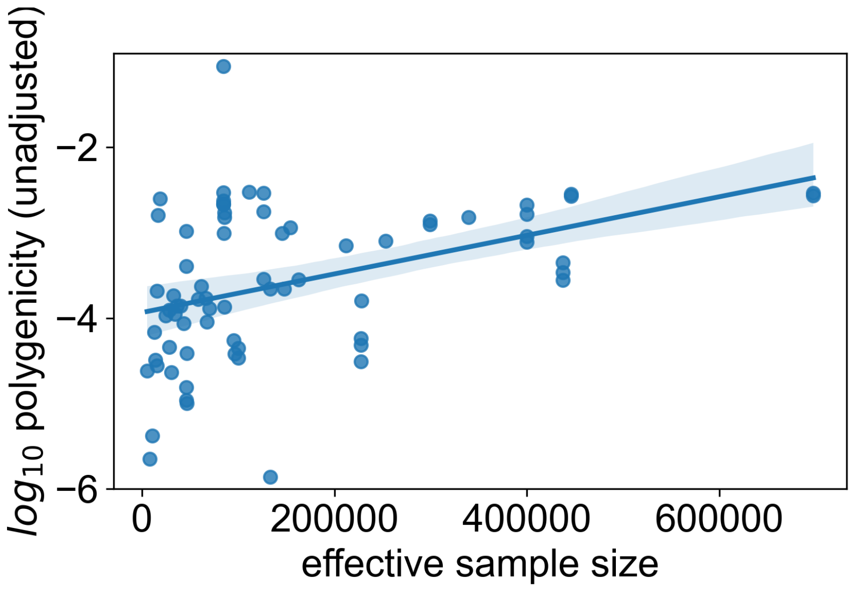
